# An RNA-centric global view of *Clostridioides difficile* reveals broad activity of Hfq in a clinically important Gram-positive bacterium

**DOI:** 10.1101/2020.08.10.244764

**Authors:** Manuela Fuchs, Vanessa Lamm-Schmidt, Falk Ponath, Laura Jenniches, Lars Barquist, Jörg Vogel, Franziska Faber

**Author notes:** authors contributed equally (Manuela Fuchs, Vanessa Lamm-Schmidt). Correspondence: Franziska Faber.

## Abstract

The Gram-positive human pathogen *Clostridioides difficile* has emerged as the leading cause of antibiotic-associated diarrhea. Despite growing evidence for a role of Hfq in RNA-based gene regulation in *C. difficile*, little is known about the bacterium’s transcriptome architecture and mechanisms of post-transcriptional control. Here, we have applied a suite of RNA-centric techniques, including transcription start site mapping, transcription termination mapping and Hfq RIP-seq, to generate a single-nucleotide resolution RNA map of *C. difficile* 630. Our transcriptome annotation provides information about 5’ and 3’ untranslated regions, operon structures and non-coding regulators, including 42 sRNAs. These transcriptome data are accessible via an open-access browser called ‘Clost-Base’. Our results indicate functionality of many conserved riboswitches and predict novel *cis*-regulatory elements upstream of MDR-type ABC transporters and transcriptional regulators. Recent studies have revealed a role of sRNA-based regulation in several Gram-positive bacteria but their involvement with the RNA-binding protein Hfq remains controversial. Here, sequencing the RNA ligands of Hfq reveals *in vivo* association of many sRNAs along with hundreds of potential target mRNAs in *C. difficile* providing evidence for a global role of Hfq in post-transcriptional regulation in a Gram-positive bacterium. Through integration of Hfq-bound transcripts and computational approaches we predict regulated target mRNAs for the novel sRNA AtcS encoding several adhesins and the conserved oligopeptide transporter *oppB* that influences sporulation initiation in *C. difficile*. Overall, these findings provide a potential mechanistic explanation for increased biofilm formation and sporulation in an *hfq* deletion strain and lay the foundation for understanding clostridial ribo regulation with implications for the infection process.

## INTRODUCTION

Antibiotic-resistant bacteria are one of the major global threats to human health, endangering our ability to perform a range of modern medical interventions. The obligate anaerobe, spore-forming *Clostridioides difficile* (*C. difficile*) has become the leading cause of antibiotic-associated diarrhea over the past two decades (Rupnik et al., 2009). In addition, *C. difficile* shows increasing numbers of multi-resistant clinical isolates (Peng et al., 2017) and recurrent infections after antibiotic therapy, which often leaves fecal microbiota transfer as the only clinical option (Khanna and Gerding, 2019).

The clinical challenges posed by *C. difficile* have prompted much effort to understand how this pathogen regulates virulence in response to environmental conditions. As a result, there is a comprehensive body of literature focusing on toxin production and sporulation control by several global metabolic regulators including CcpA, CodY, Rex and PrdR (Bouillaut et al., 2015; Martin-Verstraete et al., 2016). Furthermore, several specialized and general sigma factors including TcdR (Martin-Verstraete et al., 2016), Spo0A (Pettit et al., 2014), SigD (El Meouche et al., 2013; McKee et al., 2013), SigH (Saujet et al., 2011) and SigB (Kint et al., 2017) have been linked to virulence and metabolism, although the exact molecular mechanisms often remain unknown. Importantly, most of this knowledge has been accumulated through detailed studies of individual genes and promoters. By contrast, RNA-seq based annotations of the global transcriptome architecture, which have accelerated research in the Gram-positive pathogens *Listeria* (Wurtzel et al., 2012), *Staphylococcus* (Mader et al., 2016) and *Streptococcus* (Warrier et al., 2018), have not been available for *C. difficile* so far.

This paucity of global knowledge about RNA output in *C. difficile* readily extends to post transcriptional control of gene expression. The bacterium is of particular scientific interest, being the only Gram-positive species thus far in which deletion of *hfq* seems to have a large impact on gene expression and bacterial physiology (Boudry et al., 2014; Caillet et al., 2014). Specifically, deletion of *hfq* increases sporulation (Maikova et al., 2019), a crucial pathogenic feature of this bacterium that enables transmission between hosts. In Gram-negative bacteria, Hfq commonly exerts global post-transcriptional control by facilitating short base pairing interactions of small regulatory RNAs (sRNAs) with *trans*-encoded target mRNAs (Holmqvist and Vogel, 2018; Kavita et al., 2018) but its role in Gram-positive bacteria remains somewhat controversial. For example, recent work on the function of Hfq in *Bacillus subtilis* (*B. subtilis)*, the model bacterium for Firmicutes, revealed *in vivo* association with a subset of sRNA (Dambach et al., 2013), but comparative analyses of a wild-type and hfq knockout strain revealed only moderate effects on sRNA and mRNA transcript levels (Hammerle et al., 2014) and the absence of any significant growth defect (Rochat et al., 2015) which lead to the conclusion that Hfq plays a minor role in post-transcriptional regulation in *B. subtilis*.

Previous efforts using *in silico* methods (Chen et al., 2011) and RNA-sequencing (Soutourina et al., 2013) have predicted >100 sRNA candidates in *C. difficile*, which would suggest the existence of a large post-transcriptional network. However, under which conditions these sRNAs are expressed, which targets they regulate, and whether they depend on Hfq remain fundamental open questions.

Another important feature of post-transcriptional control in *C. difficile* are *cis*-regulatory RNA elements. There has been pioneering work deciphering the function of genetic switches located in the 5’ untranslated region (5’ UTR) of the *flgB* operon, containing the early stage flagellar genes, and the cell wall protein encoding gene *cwpV* (Anjuwon-Foster and Tamayo, 2017; Emerson et al., 2009). Moreover, cyclic di-GMP responsive riboswitches were shown to regulate biofilm formation and toxin production (Bordeleau et al., 2011; McKee et al., 2018; McKee et al., 2013; Peltier et al., 2015). However, other members of the many different riboswitch classes or common *cis*-regulatory elements such as RNA thermometers have not been systematically searched for. Combined with the nascent stage of sRNA biology in *C. difficile*, this argues that global approaches are needed to understand the full scope of post-transcriptional regulation in this important human pathogen.

In the present study, we have applied recently developed methods of bacterial RNA biology (Hor et al., 2018) to construct a global atlas of transcriptional and post-transcriptional control in *C. difficile*. Our approach demonstrates how integration of different RNA-sequencing based methods readily reveals new regulatory functions. Accordingly, we were able to predict regulatory interactions between a newly annotated Hfq-binding sRNA and target mRNA candidates. Overall, we provide evidence for extensive Hfq-dependent post-transcriptional regulation and provide the foundation for future mechanistic studies of RNA-based gene regulation in *C. difficile*.

## RESULTS

### High-resolution transcriptome maps of C. difficile 630

For generating high-resolution transcriptome maps of *C. difficile*, we chose the toxigenic reference strain 630 (DSM 27543, GenBank: CP010905.2). Being widely used by the *C. difficile* community, this strain offers the most comprehensive genome annotation. Using two different global RNA-seq approaches, we analyzed RNA samples from three different conditions: late-exponential and early-stationary phase of growth in Tryptone-yeast broth, as well as late-exponential phase in Brain-heart-infusion broth (Fig. 1A). The resulting genome-wide maps provide single-nucleotide resolution transcriptional start sites (TSSs), transcript ends, 5’ and 3’ untranslated regions (UTRs) and operon structures (Fig. 1B). In addition, we have used this information to correct previous ORF annotations, to add previously overlooked small genes and to annotate sRNA loci. Inspired by other online gene expression databases such as SalCom (Kroger et al., 2013), AcinetoCom (Kröger et al., 2018) and Theta-Base (Ryan et al., 2020), we have launched an open-access interactive web browser, called ‘Clost-Base’ (https://www.helmholtz-hiri.de/en/datasets/clostridium), for our transcriptome data. This online resource for *C. difficile* allows easy visualization of the transcriptomic data in the context of annotated coding and non-coding genes as well as transcript features (e.g. transcription start sites and transcription termination sites) that we have experimentally determined in this work. The browser enables search queries and retrieval of primary sequences for any annotated feature in the database and is a valuable resource for the *C. difficile* community.

**Figure 1.**
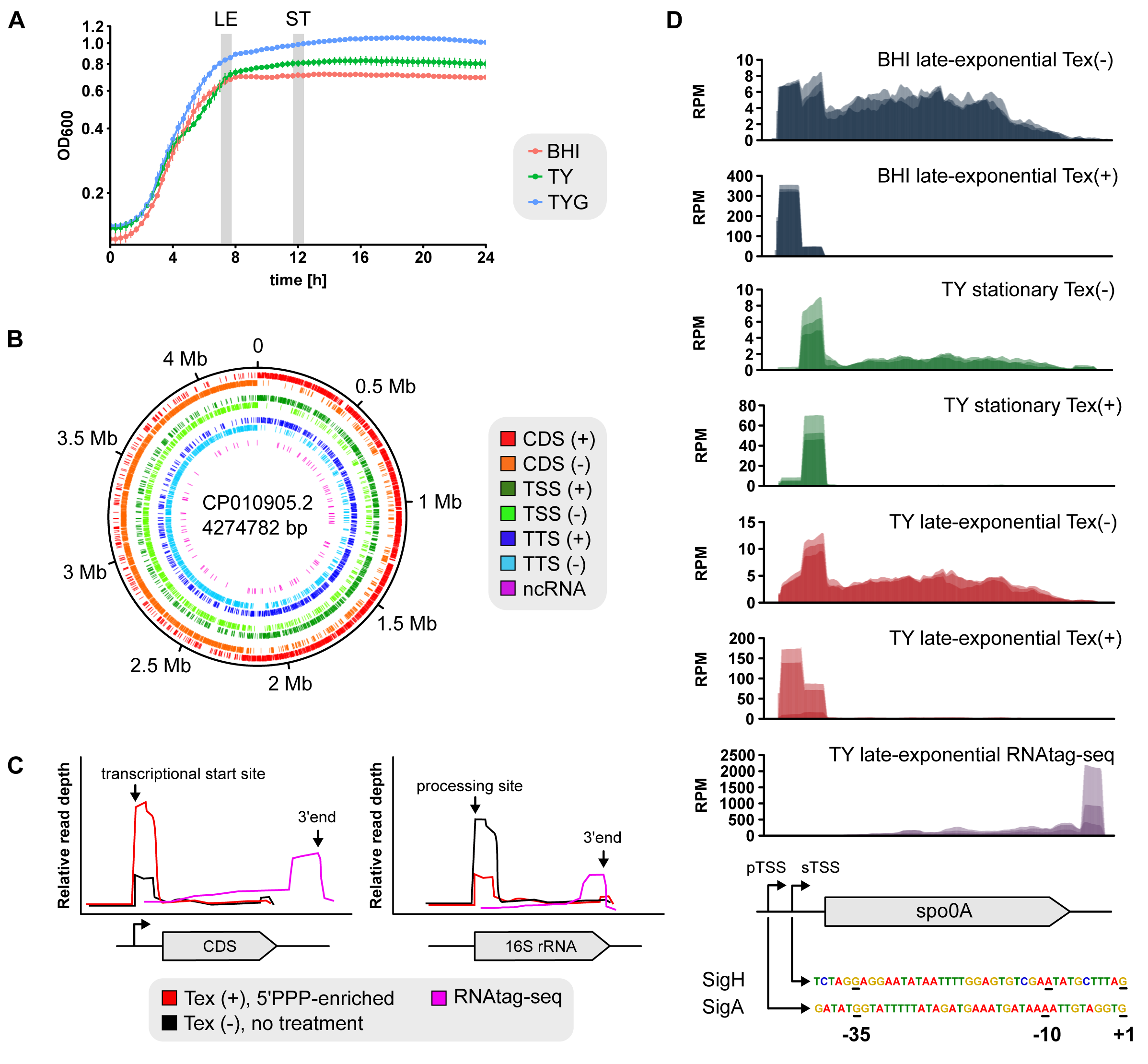
Global RNA-seq approaches for high-resolution transcriptome mapping. (A) Sequencing samples were generated from strain *C. difficile* 630 grown to late-exponential and stationary phase of growth in Tryptone-yeast broth, as well as late-exponential phase in Brain-heart-infusion broth. (B) Distribution of newly annotated features (TSSs, TTSs and non-coding RNAs) across the genome of *C. difficile* strain 630. (C) Read profiles generated with dRNA-seq and RNAtag-Seq allow the annotation of transcriptional start sites (TSSs) and transcriptional termination sites (TTSs). For dRNA-seq one fraction of total RNA is treated with terminator exonuclease (TEX+), which specifically degrades processed transcripts carrying a 5′ monophosphate. The other fraction remains untreated. This differential treatment results in a relative read enrichment for primary transcripts in the TEX+ treated libraries allowing the identification of TSSs. For RNAtag-Seq adapters for sequencing are ligated to RNA 3’ ends thereby capturing 3’ ends of transcripts. (D) Benchmarking of dRNAseq approach. Two experimentally determined growth-phase dependent TSS for *spo0A* are consistent with dRNA-seq based identification of *spo0A* associated TSSs. Abbreviations: LE, late-exponential growth phase; ST, stationary growth phase; CDS, coding sequence; TSS, transcriptional start site; TTS, transcript termination site.

### Genome-wide annotation of transcription start sites

Differential RNA-seq (dRNA-seq) (Sharma et al., 2010; Sharma and Vogel, 2014) was performed to capture 5’ ends of transcripts, a method relying on the differential treatment of input RNA sample with terminator exonuclease (Tex). In brief, one half of the sample remains untreated (Tex-) to capture both primary (5′- PPP) and processed (5′-P or 5′OH) 5’ ends of transcripts; the other half is treated (Tex+) leading to the specific degradation of processed (5′-P or 5′OH) RNAs, thus enriching primary transcripts and enabling TTS annotation (Fig. 1C). Conversely, relative read enrichment in the Tex− cDNA libraries indicate RNA processing sites (Fig. 1C). With a genome size of 4,274,782 bp, the *C. difficile* 630 genome comprises annotations for 3,778 predicted coding sequences as well as tRNAs, rRNAs and the housekeeping RNAs 6S RNA, RNaseP, tmRNA and SRP. By identifying 2,293 transcriptional start sites (TSS), we were able to define transcriptional units of individual genes and polycistronic operons for approximately half of the genome (Table S1).

Benchmarking our global data with previous studies of individual genes (Emerson et al., 2009; Saujet et al., 2011; Wydau-Dematteis et al., 2018), our dRNA-seq results are fully consistent with published TSSs of *sigH*, *cwpV*, *cwp19*, and *spo0A* (Fig. 1D). For annotation purposes, we assigned TSS to one of five classes according to their genomic location and expression level: pTSS (primary TSS of a gene or operon), sTSS (secondary TSS showing lower expression level compared to pTSS for the same gene or operon), iTSS (internal TSS located inside a gene), aTSS (antisense TSS to a gene within 100 nt distance) and oTSS (orphan TSS, no nearby gene) (Fig. 2A). Naturally, some of these TSS annotations overlap, for example, among the 1627 pTSS, 126 are located within a gene (iTSS) and 46 are transcribed antisense (aTSS) to a gene. However, in contrast to other bacteria where antisense transcription is a pervasive transcriptome feature (Wade and Grainger, 2014), it only accounts for approx. 6% of TSS events in *C. difficile*.

**Figure 2.**
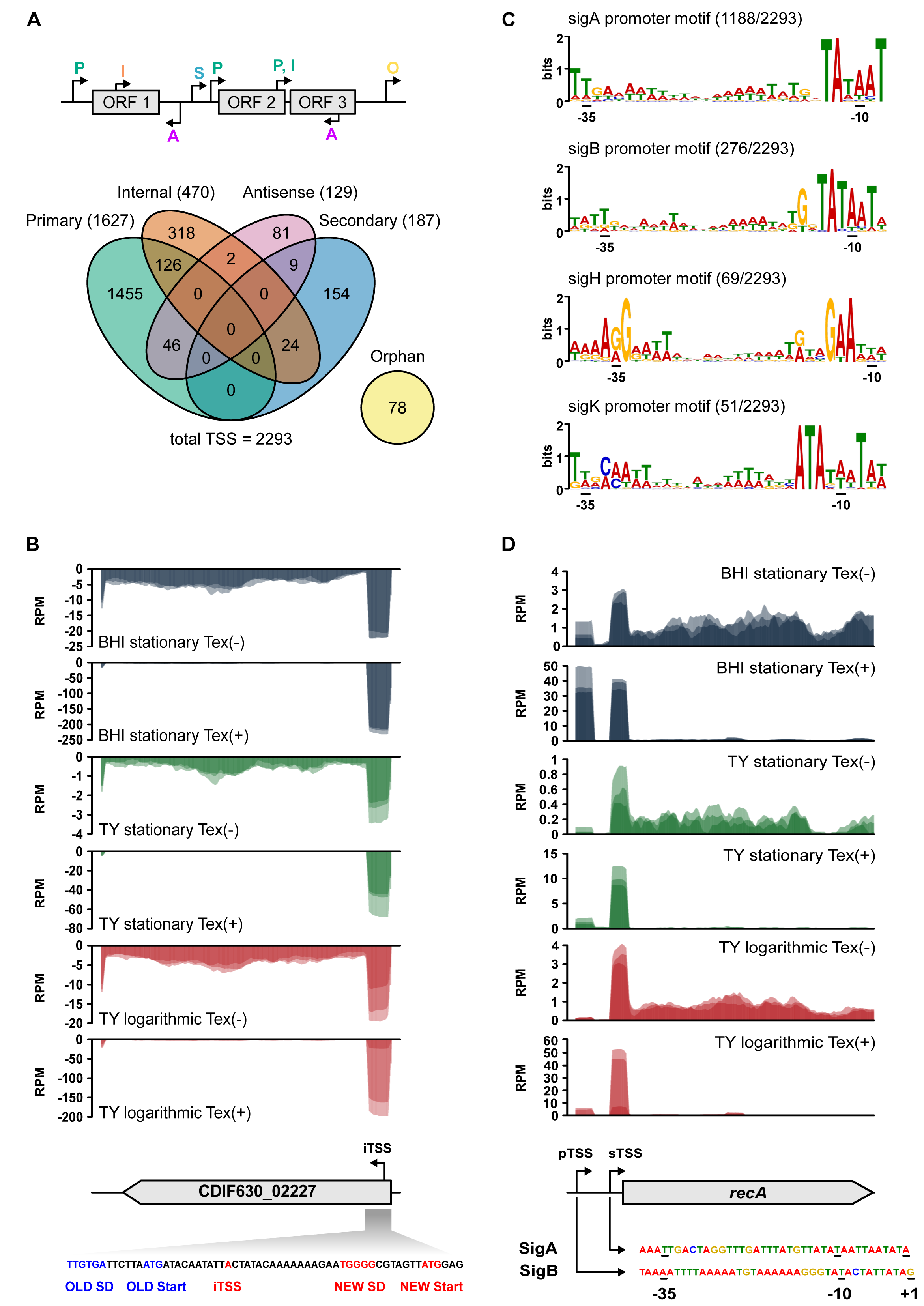
Features associated with transcriptional start sites. (A) Top: TSS classification based on expression strength and genomic location: primary (P), secondary (S), internal (I), antisense (A) and orphan (O). Bottom: Venn diagram showing distribution of TSSs among classes. (B) Re-annotation of CDIF630_02227 gene based on a newly identified internal TSS 30 bases downstream of its annotated start codon. Reads of dRNA-seq libraries (TEX+/TEX-) mapping to the CDIF630_02227 region are shown above the sequence covering the region around the old and new start codon. The old and new AUGs and associated Shine-Dalgarno (SD) sequences are indicated. (C) Promoter regions associated with detected TSSs were analysed using the MEME suite. Promoter motifs were recovered for SigA, SigH, SigB and SigK. (D) Two alternative promoter sequences for SigA and SigB are associated with a pTSS and sTSS of *recA*, respectively.

Internal TSS were abundant and accounted for 20% of all start sites, many of which seem to uncouple downstream genes within operons. However, they may also indicate mis-annotated translational start codons. One example is a putative hydrolase (CDIF630_02227) encoded between *ermB1* and *ermB2* of the erythromycin resistance cassette. We detected an iTSS located 14 bases downstream of the presently annotated canonical start codon. However, we manually checked for canonical (AUG) or alternative start codons (UUG, GUG) downstream of the annotated start codon, including a ribosome-binding site (RBS). This approach resulted in the re-annotation of the CDIF630_02227 ORF with the new canonical start codon 43 bases downstream of the existing ORF annotation, placing a RBS in optimal distance to the new start codon and generating a 5’ UTR length of 29 nucleotides (Fig. 2B). Similarly, we propose re-annotations in eight additional cases, including the *spaFEGRK* operon encoding an antibiotic/multidrug-family ABC transport system (Table S2).

### Promoter architectures

*C. difficile* encodes a high number of σ factors which serve specialized functions, such as the regulation of toxin production or sporulation. Consensus sequences have been proposed for several of them including the vegetative SigA (Soutourina et al., 2013), the general stress response SigB (Kint et al., 2019; Kint et al., 2017), the major transition phase SigH (Saujet et al., 2011) and the sporulation-specific SigK (Saujet et al., 2013) sigma factors, mostly based on comparative transcriptome analyses of respective deletion strains and their wild-type strain. By applying MEME-based searches upstream of all TSS we were able to refine published consensus sequences for all these four σ factors (Fig. 2C) and substantially extended their network of regulated genes (Table S3 and S1).

Unsurprisingly, approximately half of the detected TSS (1,188/2,293) were associated with a SigA-type promoter (Table S3), which include the previously determined TSS of *sigH* and transcriptional regulator gene *clnR* (Saujet et al., 2011; Serrano et al., 2016; Woods et al., 2018). The general stress factor SigB is another widespread sigma factor in gram-positive bacteria (van Schaik and Abee, 2005). However, among Clostridia, *C. difficile* is the only species encoding an annotated homologue. At the onset of stationary phase, SigB controls about 25% of all *C. difficile* genes, primarily those involved in metabolism, sporulation and stress responses (Kint et al., 2017). We were able to extend the number of experimentally mapped SigB-associated TSS from 20 (Kint et al., 2017) to 277 (Table S3). Functional categories assigned to those genes are DNA integration and recombination, transcriptional regulation, and cell wall turnover. Similarly, we extend the SigH regulon to 69 genes, which previously included approx. 40 genes or operons, respectively (Saujet et al., 2011). SigH is the key σ factor of transition phase and sporulation initiation in *C. difficile*. Many genes associated with a SigH-type promoter in our analysis have functions related to the biosynthesis of amino acids, secondary metabolites and antibiotics, in addition to general metabolic pathways. Further, we identified 52 genes associated with a promoter signature for sporulation-specific sigma factor SigK, including known genes of the SigK regulon such as *sleC* and *cdeC* (Pishdadian et al., 2015). In addition, we found 39 genes with alternative TSSs that were associated with two different promoter sequences. For example, RecA, which controls the DNA damage response by homologous recombinational repair of damaged DNA, is associated with a SigA- and SigB-type promoter supporting its role in stress responses (Fig. 2D).

### Global mapping of transcript ends

To map transcript 3’ termini, we adopted the RNAtag-Seq protocol (Shishkin et al., 2015), a technique utilizing initial adapter ligation to exposed RNA 3’ ends. This approach yielded 2,042 experimentally determined transcript 3’ ends, which were assigned to one of the following classes according to their genomic location: 3’ UTR (downstream of an annotated CDS or non-coding RNA locus), 5’ UTR (between the TSS and the start codon of an annotated CDS), CDS (within a coding sequence), orphan (downstream of an orphan TSS) and CRISPR (associated with CRISPR array) (Fig. 3A). The majority of transcription termination events mapped to the region downstream of annotated genes allowing us to generate a genome-wide map of 3’ UTRs (Table S4). Our data reveal that 42% of all detected *C. difficile* 3’ UTRs downstream of an annotated CDS are >100 bp in length (Fig. 3C), which resembles the 3’ UTR length distribution in *Staphylococcus aureus* (Ruiz de los Mozos et al., 2013).

**Figure 3.**
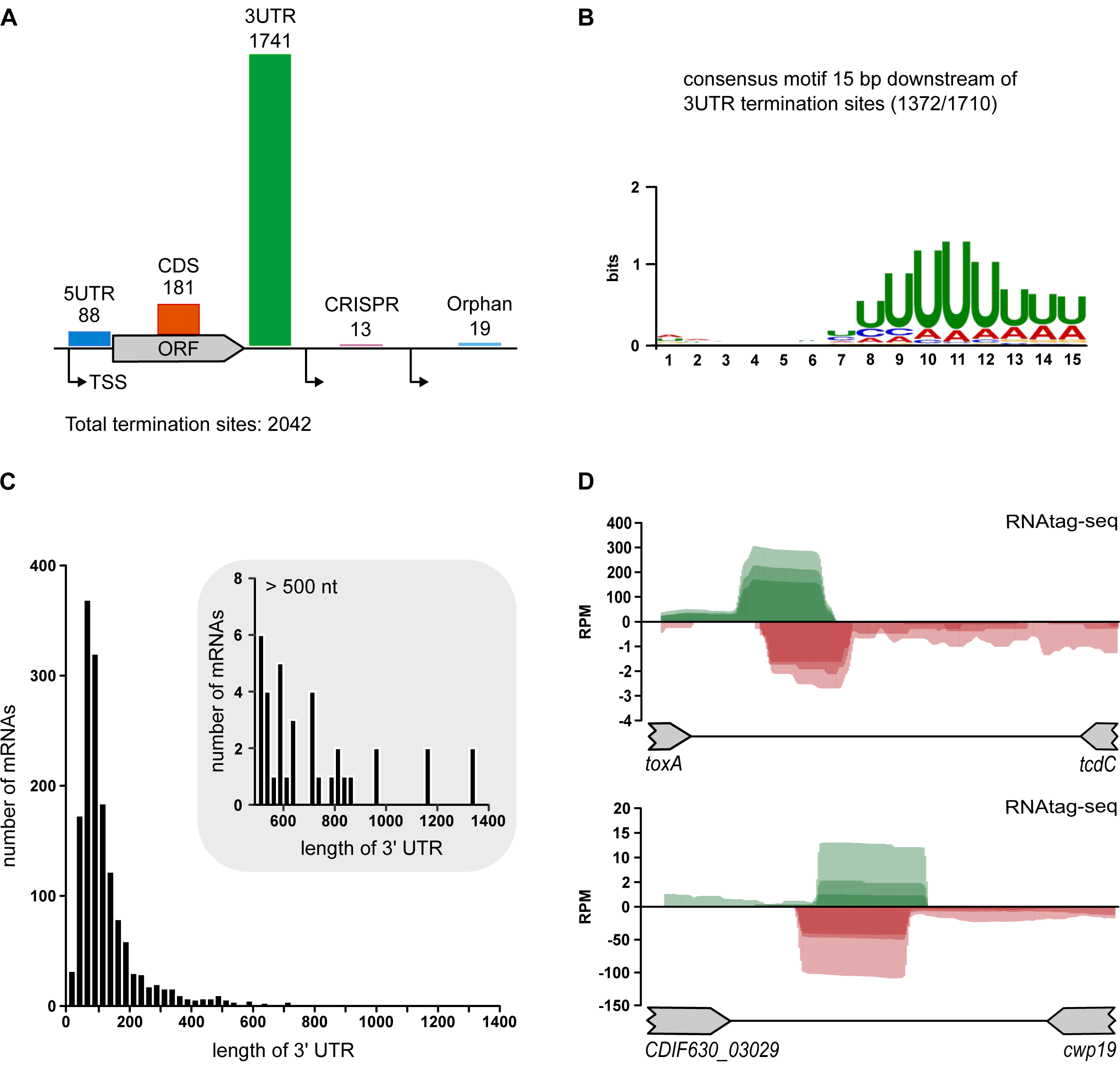
Features associated with transcriptional termination sites. (A) Top: TTS classification based on genomic location: 3UTR (downstream of CDS or non-coding gene), 5UTR (within the 5’ UTR of a coding sequence), CDS (within coding sequence), orphan (downstream of orphan TSS), CRISPR (associated with CRISPR array). (B) Frequencies of 3’ UTR lengths based on 1741 TTSs. (C) Consensus motif associated with 3’ UTR termination sites was identified using the MEME suite. (D) Reads in the RNAtag-Seq libraries reveal overlapping transcription termination sites for convergently transcribed gene pairs.

For the majority of 3’ UTR termination sites (75%) we identified a consensus motif that is in agreement with known sequence features of Rho-independent terminators (Fig. 3C and Table S4). This indicates a major role of Rho-independent termination in *C. difficile* and resembles transcription termination in *B. subtilis* where the terminator protein Rho is not essential, regulating transcription termination for only a few genes including *rho* itself (Ingham et al., 1999; Nicolas et al., 2012; Quirk et al., 1993). Further, recent studies in *Escherichia coli* (*E. coli*) showed that bidirectional transcription termination is a pervasive transcriptome feature in bacteria (Ju et al., 2019). Accordingly, our analysis of transcription termination sites in *C. difficile* revealed many overlapping termination events located between convergently transcribed genes, such as for *cwp19* and its convergent gene CDIF630_03029 or for the toxin gene *toxA* and the negative regulator of toxin gene expression *tcdC* (Fig. 3D).

### Small ORFs

High-resolution transcriptome maps allow the identification of ORFs that have been overlooked in automated genome annotation. This includes so called small ORFs (sORFs) of usually 50 amino acids or less, an emerging class of bacterial genes with an unfolding spectrum of new biological functions (Garai and Blanc-Potard, 2020; Hemm et al., 2020). In many cases, they are predicted to be membrane proteins, containing an alpha-helical transmembrane domain (Storz et al., 2014). Focusing on oTSS in particular, we searched for novel ORFs based on the following criteria (i) presence of a start and stop codon (ii) presence of a RBS within 15 bp upstream of the start codon and (iii) sequence conservation in other *Clostridioides* strains. Based on this approach, we identified 12 sORF candidates, seven of which are predicted to contain a transmembrane helix (Table S2). Among the identified candidates six are toxins that are part of previously identified *C. difficile* type-I toxin-antitoxins systems which were not annotated (Maikova et al., 2018; Soutourina, 2019). Among the remaining sORFs four are high confidence candidates; one being a conjugal transfer protein with annotation in other *C. difficile* strains; two candidates that each have a Shine-Dalgarno (SD) sequence and a predicted α-helical transmembrane domain; and one sORF associated with a 160nt long 5’ UTR region harboring a c-di-GMP-I riboswitch. This association with a cyclic di-GMP responsive riboswitch suggests a potential virulence-associated function since c-di-GMP regulates not only motility and biofilm formation, but also toxin production (Bordeleau et al., 2011; McKee et al., 2018; McKee et al., 2013; Peltier et al., 2015).

### 5’ UTRs and associated regulatory elements

Bacterial 5’ UTRs can influence gene expression in response to environmental signals, usually through embedded *cis*-regulatory elements such as riboswitches, RNA thermometers and genetic switches. Similar to other bacterial species, such as *B. subtilis* and *Listeria monocytogenes*, the majority of 5’ UTRs in *C. difficile* range from 20 to 60 nucleotides in length (Fig. 4A, Table S1; (Irnov et al., 2010; Sharma et al., 2010; Wurtzel et al., 2012). An aGGAGg motif that likely serves as an RBS was detected in ~90% of these experimentally mapped 5’ UTRs (Fig. 4A, inlet). In addition, we identified six leaderless mRNAs with a 5’ UTR of <10 nucleotides in length, including the *spoVAE* gene within the tricistronic *spoVACDE* operon (Donnelly et al., 2016) indicating a potential transcription of *spoVAE* that is uncoupled from the other two genes of the proposed operon.

**Figure 4.**
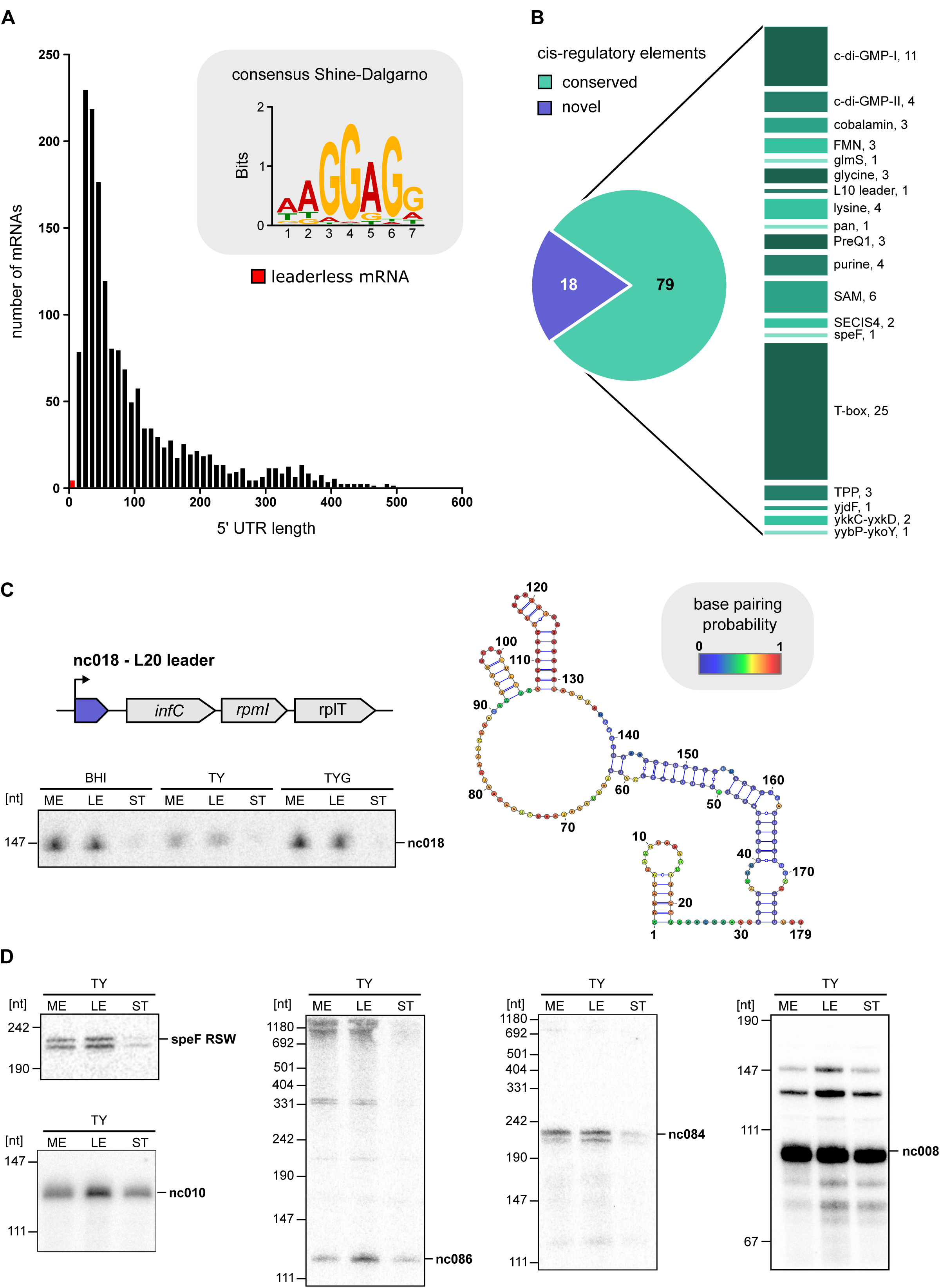
Discovery of *cis*-regulatory elements in 5’ UTR regions. (A) Frequencies of 5’ UTR lengths based on 1646 primary and secondary TSSs. Red bars indicate six leaderless mRNAs with a 5’ UTR lengths of <10 nt. The inset shows the SD sequence motif of *C. difficile* 630. (B) Classification of premature transcription termination (PTT) events in 5’ UTRs of mRNAs. The majority is associated with known RNA families. The remaining 19 PTT events lack homology to known riboregulators and are putative novel regulators. (C) Putative L20 leader in the 5’ UTR of the *infC* operon. Left: schematic overview of the *infC* operon organization including 5’ UTR region and expression analysis of CDIF630nc_00018 by Northern blot. Right: Predicted secondary structure for CDIF630nc_00018 using RNAfold. Nucleotides are colored according to base-pairing probabilities. (D) Expression analysis of novel 5’ UTR *cis*-regulatory elements. Total RNA was extracted at mid-exponential, late-exponential and stationary growth phases from *C. difficile* 630 grown in TY medium and analyzed by northern blot using radioactively labeled DNA probes.

In addition to the average sized 5’ UTRs, a high number of genes are associated with surprisingly long 5’ UTRs: 561 possess a 5’ UTR of >100 nucleotides, and 129 of those have a very long 5’ UTR of >300 nucleotides. The latter include the 496 nt long 5’ UTR of *flgB*, which was previously shown to contain both a c-di GMP-I riboswitch and a genetic switch that co-regulate the expression of *C. difficile* flagella and toxin genes (Anjuwon-Foster and Tamayo, 2017). Further analysis of these 5’ UTRs resulted in the identification of 77 Rfam-predicted riboswitch candidates (Kalvari et al., 2018) (Fig. 4B). According to our RNAtag-Seq data, premature transcription termination was evident in 57 of those 77 predicted riboswitches, indicating their ON state in which transcription of the parental gene is repressed (Table S5). Interestingly, we detected a *speF* riboswitch previously only documented in gram-negative alpha-proteobacteria associated with CDIF630_01955, encoding a methyltransferase domain protein. As before, the associated termination site (ON state) suggests that the riboswitch is functional, which is further supported by a 220 nt northern blot signal in samples extracted from cultures grown in three different media (Fig. 4D).

Additionally, we found two putative c-di GMP riboswitches to be associated with oTSSs: a c-di GMP-I riboswitch in the vicinity of and opposite to a CRISPR12 array, associated with a predicted sORF (see section New ORFs and Table S5) and a c-di GMP-II riboswitch immediately downstream of *prdC*, encoding a subunit of the proline reductase enzyme. The latter is >900 nt away from the closest coding region. No putative sORF was identified in this case, instead, the associated regulated transcript might be an sRNA as published for three cobalamin riboswitches in *Enterococcus faecalis*, *Listeria monocytogenes* and *Streptococcus sanguinis* each regulating an sRNA (DebRoy et al., 2014; Mellin et al., 2014).

Besides riboswitch-associated premature termination, we observed 17 additional events of putative premature transcription termination (PTT) in 5’ UTRs that lack similarity to conserved riboregulators. A potential generation from mRNA processing is unlikely in these cases, since the characteristic read enrichment in the untreated (TEX-) cDNA library, usually associated with processing sites, is missing. One such PTT event is located in the 5’ UTR of the *infC*-*rpmI*-*rplT* operon whose expression is controlled through an L20 leader via a transcription attenuation mechanism in *B. subtilis* (Babina et al., 2018; Choonee et al., 2007). This ribosomal protein leader autoregulatory structure is also found in other low-GC gram-positive bacteria (Rfam family RF00558) but does not seem to be conserved on the primary sequence level in *C. difficile*. Nevertheless, secondary structure prediction reveals extensive interactions between distant bases that is reminiscent of the antiterminator conformation of the L20 leader from *B. subtilis* (Fig. 4C).

Further genes associated with such PTT events include several PTS systems (*bglF*, *bglF1*, *bglG3* and *bglG4*) that are known to be regulated by antiterminator proteins in *B. subtilis* (Fujita, 2009). Further, we also detected PTT events within the 5’ UTR of genes encoding transcriptional regulators (CDIF630_00097, CDIF630_02384, CDIF630_02922), MDR-type ABC transporters (CDIF630_03083, CDIF630_02847, CDIF630_03664) and *aroF* (Table S5). Northern blot validation for a selection of candidates confirmed the RNA-seq predicted transcript sizes and revealed larger bands in several cases that are likely to correspond to the full-length parental gene or its degradation products (Fig. 4D).

### RIP-seq identifies Hfq as a global RNA-binding protein in C. difficile

Our primary transcriptome analysis identified 42 novel transcripts that lack an internal open reading frame, qualifying them as potential small regulatory RNAs (sRNA) (listed in Table S5). A classification based on their genomic location (Fig. 5A) revealed the largest group to be 3’ UTR derived sRNAs (18), followed by those encoded cis-antisense either to a gene, another sRNA or the 5’/3’ UTRs of coding sequences (13). In comparison, only few sRNAs were located in intergenic regions (8) or derived from the 5’ UTR of mRNAs (3). Except for two transposon associated sRNAs (CDIF630nc_00004 and CDIF630nc_00069) and one located on a prophage (CDIF630nc_00095), all of them are highly conserved among *C. difficile* strains, whereas sequence conservation beyond *C. difficile* was extremely rare (Fig. S1, Table S7).

**Figure 5.**
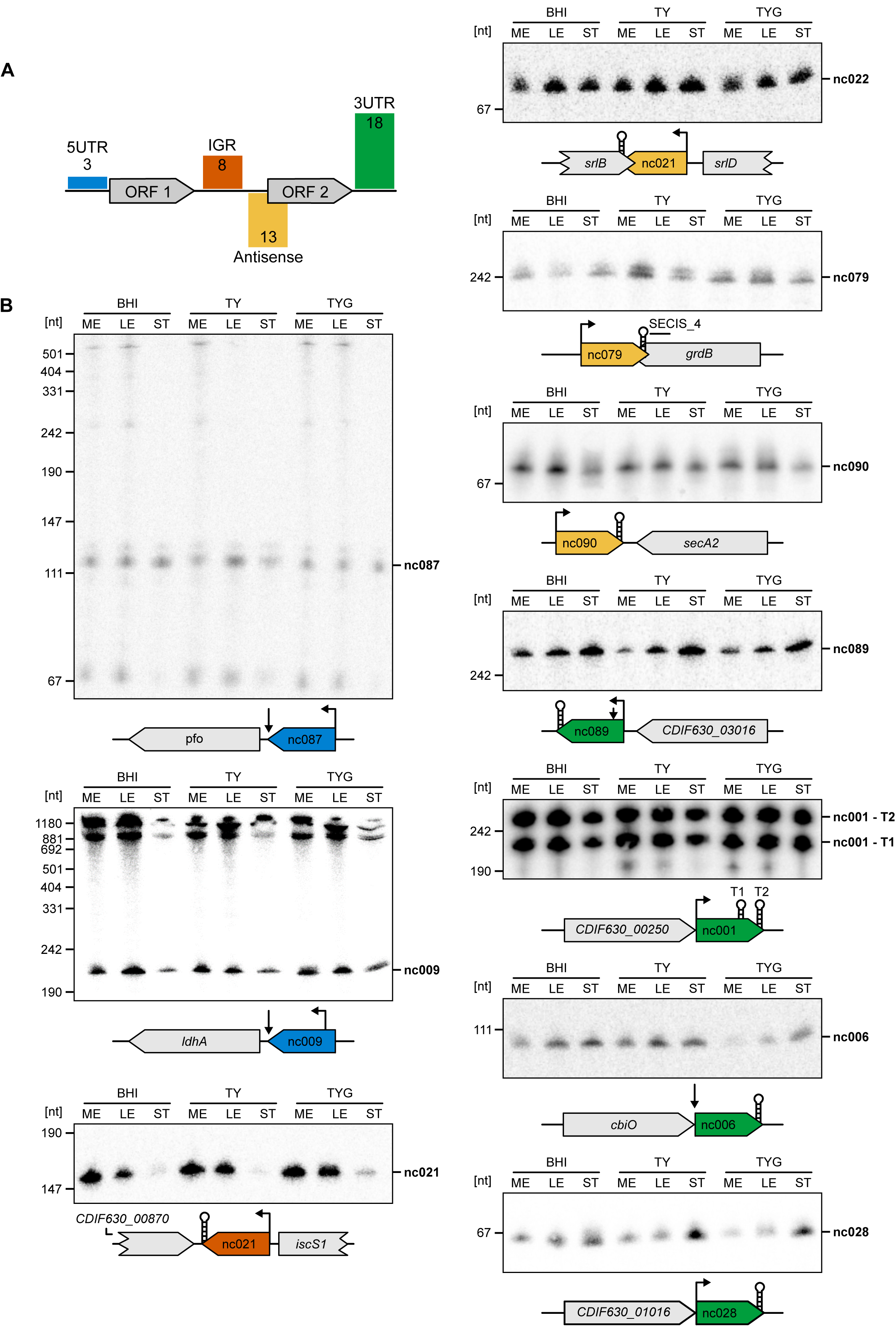
The sRNA landscape of *C. difficile* 630. (A) Classification of annotated sRNA candidates based on their genomic location. Numbers indicate amount of annotated sRNA candidates for each class (B) Expression profiles of representative candidates for each sRNA class. Total RNA was extracted at mid-exponential, late-exponential and stationary growth phases from *C. difficile* 630 grown in TY, TYG and BHI medium and analyzed by northern blot using radioactively labeled DNA probes.

Northern blot validation of predicted sRNA candidates revealed that that most sRNA candidates are expressed throughout exponential growth (Fig. 5B). About half of them are downregulated after entry into stationary phase, while three candidates (CDIF630nc_00028 and CDIF630nc_00089 and CDIF630nc_00105) show a marked accumulation. In agreement with the increased abundance during stationary growth phase, CDIF630nc_00028 has a predicted promoter sequences for SigB while CDIF630nc_00089 is associated with a SigH promoter signature.

In many gram-negative model organisms such as *Salmonella enterica* and *E. coli* sRNAs often require the RNA-binding protein Hfq to facilitate their interaction with target mRNAs (Hör et al., 2020). However, whether Hfq functions as a central RNA-binding protein (RBP) in *C. difficile* remains unknown. Therefore, we performed Hfq-immunoprecipitation followed by sequencing of bound RNA species (RIP-seq) in a strain expressing C-terminally 3xFLAG tagged *hfq* (Hfq-FLAG) along with its 5’ UTR under its native promoter to draft the spectrum of RNAs that are associating with Hfq *in vivo*. Bacteria were grown in TY (tryptone yeast) medium to mid-exponential, late exponential and stationary phase (Fig. 6A) and subjected to the RIP-seq protocol (Fig. 6B). Western blot validation using monoclonal FLAG antibody confirmed expression of Hfq throughout all growth phases and specific enrichment of Hfq-FLAG from *C. difficile* 630 lysate (Fig. 6C). In addition, co-purification of Hfq-bound sRNAs was confirmed by northern blot analysis of lysates (L) and eluate (E) fractions from Hfq-FLAG and Hfq-ctrl bacterial cultures (Fig. 6C).

**Figure 6.**
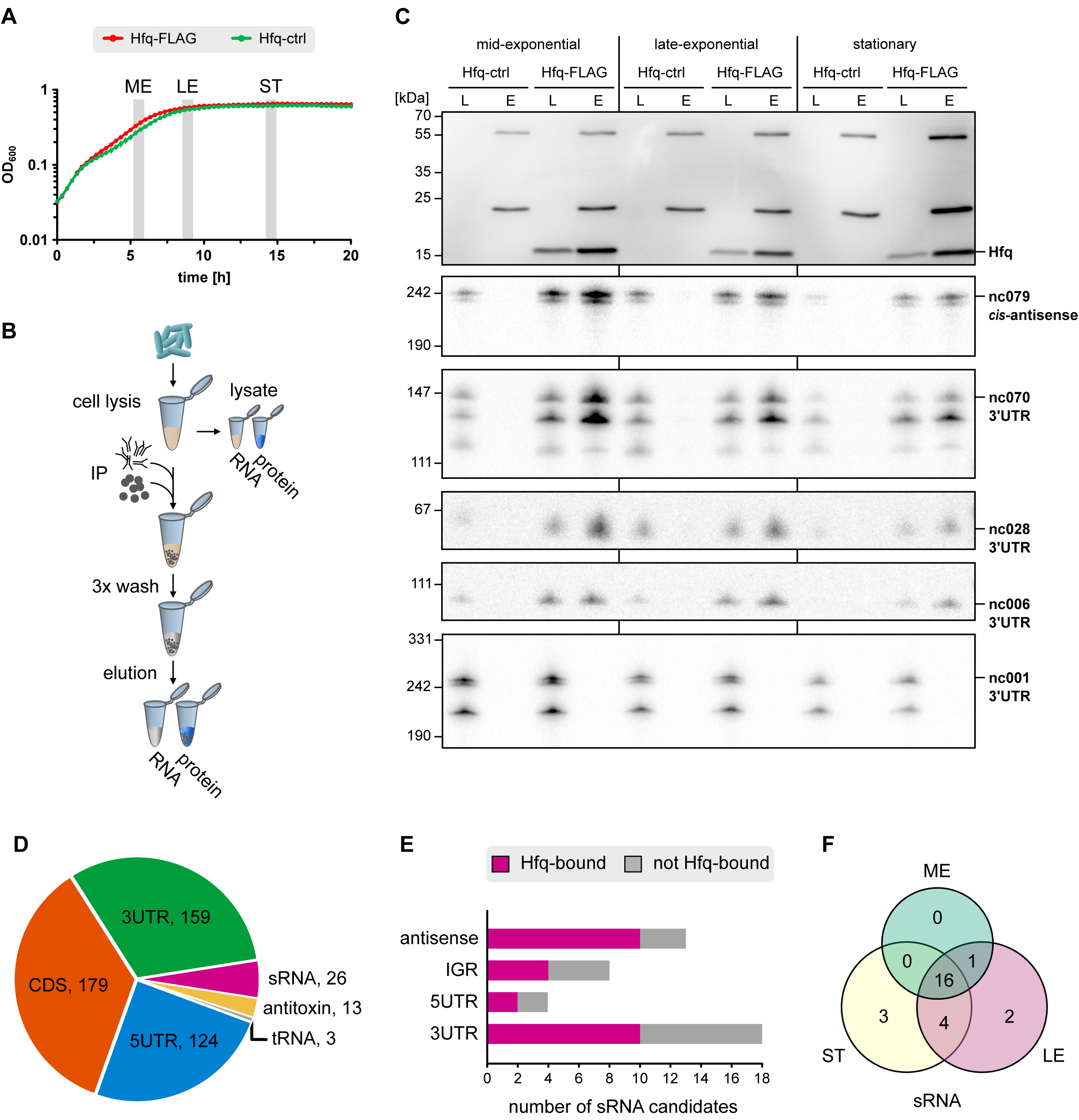
The spectrum of Hfq-associated RNA classes in *C. difficile* 630. (A) Hfq RIP-seq was performed on WT and 3xFLAG tagged hfq strains grown to mid-exponential, late-exponential and early-stationary phases in TY medium in two independent experiments. (B) Overview of RIP-seq workflow. (C) Pull-down of 3xFLAG tagged Hfq was validated by Western blot using anti-FLAG antibody. Hfq-associated RNAs were validated by northern blot using specific DNA probes. (D) Pie chart for Hfq RIP-seq showing the relative amount of Hfq-associated sequences mapping to different RNA classes. (E) Bar chart showing the fraction of sRNAs for each class that were bound by Hfq *in vivo*. (F) Number of Hfq-bound sRNA at different growth phases. Abbreviations: ME, mid-exponential; LE, late-exponential; ST, stationary; IP, immunoprecipitation; L: lysate; E: elution.

The majority of RNA species that co-immunoprecipitated with Hfq-FLAG were coding and non-coding regions of mRNAs (CDS, 5’ UTRs and 3’ UTRs) (Fig. 6D). Interestingly, we observed enrichment of many 5’ UTRs harboring riboswitches in addition to all but one type-I antitoxin transcripts (Table S6). Importantly, the majority of putative sRNAs associated with Hfq (Fig. 6E, see also Table S5). Among them was CDIF630nc_00070 (Fig. 6C), an sRNA previously shown to bind to Hfq *in vitro* and *in vivo* (Soutourina et al., 2013). Interestingly, classifying Hfq-associated sRNA with respect to their genomic location revealed that the majority of *cis*-antisense encoded sRNAs in *C. difficile* were bound by Hfq (Fig. 6E). Profiling of Hfq-bound sRNAs revealed that the majority was bound by Hfq across all three analysed growth phases with CDIF630nc_00070 and CDIF630nc_00079 being the two top-enriched sRNAs (Fig. 6F, Fig S2A and Table S6). However, a few sRNAs display a growth-phase dependent association with Hfq, such as CDIF630nc_00090 which is only enriched in stationary growth phase (Fig S2A).

### Prediction of regulatory interactions for the novel Hfq-dependent sRNA AtcS

In addition to many sRNA candidates, we observed widespread binding of mRNA transcripts (CDS, 5’ UTRs and 3’ UTRs) (Fig. 6D, Table S6) who showed a much stronger growth-phase dependent association with Hfq (Fig. 7A). In total, we identified 332 mRNAs as significantly enriched in our Hfq pull-down (log2 ≥ 2). Those mainly included genes involved in the biosynthesis of amino acids, as well as membrane transport, signal transduction and translation. Most interestingly, we also found several mRNAs encoding for proteins important for virulence in *C. difficile* such as genes involved in sporulation (*spo0A*, *spoVB*, *oppB*, *sigG* and *sspA2*), toxin production (*tcdE*), motility (*flgB*, *fliJ*, *flhB* and *cheW1*), quorum sensing (*luxS* and *agrB*), the cell wall protein encoding gene *cwpV* as well as several adhesins potentially involved in biofilm formation (CDIF630_03429, CDIF630_01459 and CDIF630_03096) (Fig. 7B, Fig. S2B, Table S6).

**Figure 7.**
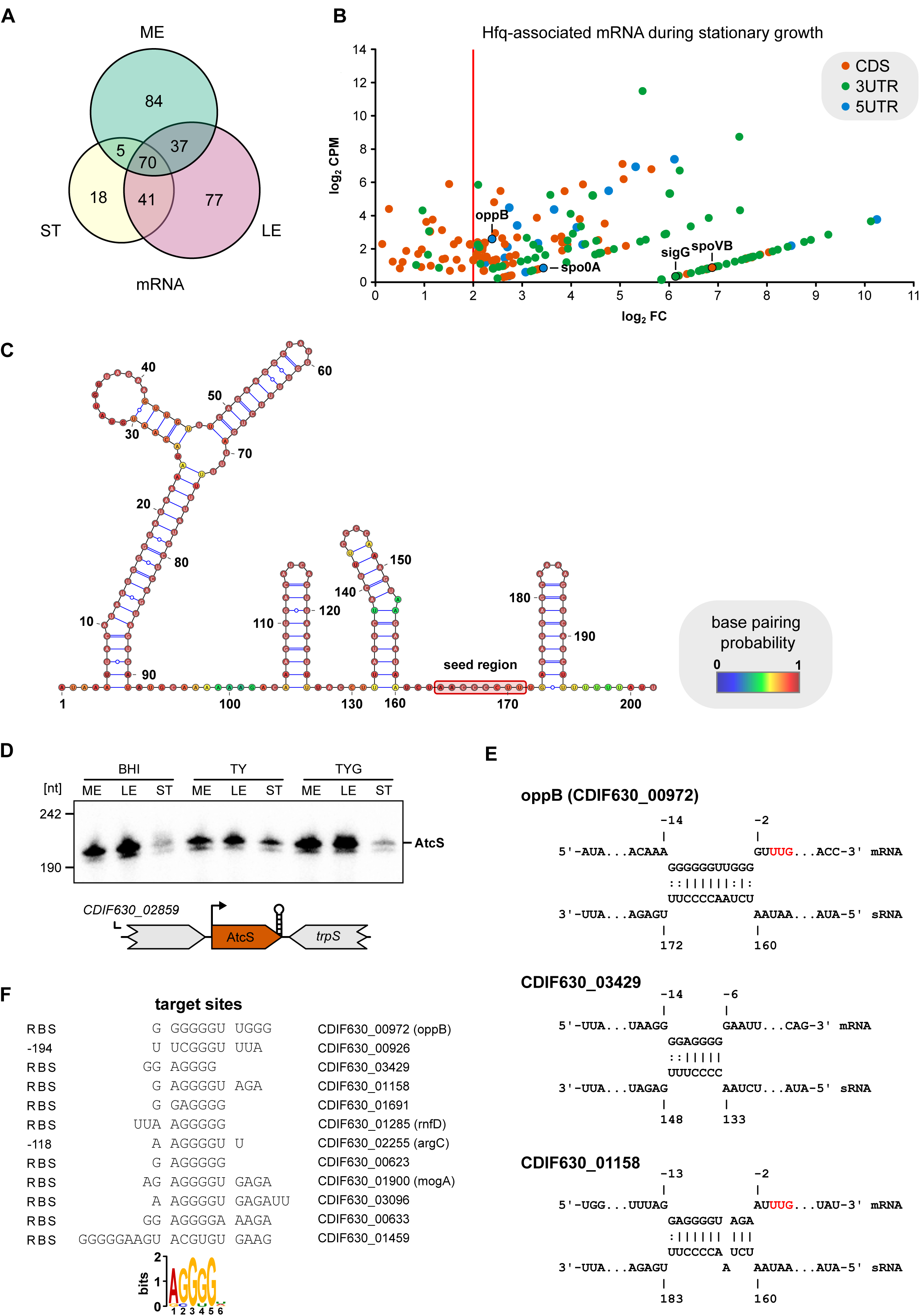
Prediction of the target spectrum for the intergenic sRNA AtcS based on Hfq-associated mRNAs. (A) Number of Hfq-bound mRNA at different growth phases. (B) Scatter-plot analysis of RIP-seq results for mRNAs that were enriched in stationary phase (log2 f.c. ≥2; cDNA read ≥10; Benjamini-Hochberg corrected P-value ≤0.1) in the 3xFLAG tagged Hfq samples. Genes encoding for sporulation-associated proteins are labelled. (C) Predicted secondary structure for AtcS using RNAfold. Nucleotides are colored according to base-pairing probabilities. Nucleotides predicted to interact with target mRNAS (seed region) are located in an accessible single-stranded stretch at the 3’ end (red box). (D) Expression profiling of AtcS by northern blot. Total RNA was extracted at mid-exponential, late-exponential and stationary growth phases from *C. difficile* 630 grown in TY, TYG and BHI medium and analyzed by northern blot using radioactively labeled DNA probes. (E) Predicted RNA duplexes formed by AtcS with selected mRNA targets. Red letters indicate the start codon. (F) MEME-based definition of a consensus motif in predicted binding sites of AtcS mRNA targets. Abbreviations: ME, mid-exponential; LE, late-exponential; ST, stationary.

To identify possible sRNA-mRNA interactions we applied the CopraRNA tool (Raden et al., 2018) to Hfq-associated sRNAs, focusing on interactions close to the RBS, since many sRNAs regulate their target mRNAs through binding close to the RBS. By comparing the *in silico* predicted potential target mRNAs with mRNAs that co-immunoprecipitated with Hfq *in vivo*, we readily identified the 202 nt *bona-fide* intergenic sRNA CDIF630nc_00085 (Fig. 7C) as having an interesting predicted target spectrum. For reasons explained below, we renamed the sRNA AtcS (adhesin-targeting *Clostridioides* sRNA).

Northern blot validation of AtcS expression indicated sustained expression during exponential growth that ceases in stationary phase, an effect that is more pronounced in glucose rich TYG medium as compared to TY and BHI (Fig. 7D). In agreement with the expression profiles, AtcS is associated with a SigA promoter motif. *In silico* secondary structure prediction revealed a moderately folded structure with an extended stem loop structure at the 5’ end, two smaller stem loops in the center of the sRNA, and the Rho-independent terminator at the 3’ end.

CopraRNA predicts at least ten mRNAs as target candidates for AtcS that were also enriched in the RIP-seq dataset (Fig. 7E&F). The corresponding predicted seed region in AtcS is located in a single-stranded stretch adjacent to the Rho-independent terminator (Fig. 7C, red circle) that base pairs with the region surrounding the SD sequence for most of the mRNA candidates. Accordingly, aligning these mRNA target sites identified the following motif: AGGGG. Targeting the region surrounding the SD sequence is a well described mechanism of sRNA-mediated regulation. Perhaps the most common outcome of this interaction is inhibition of translation by preventing ribosome binding to the mRNA (Hör et al., 2020).

Among the target candidates for AtcS, three are adhesins, of which two have homologs in *Staphylococcus epidermidis* (CDIF630_03429 and CDIF630_01459) where they are involved in surface adhesion and biofilm formation (Arrecubieta et al., 2007). Another predicted target is the oligopeptide transporter *oppB* (CDIF630_00972), encoding a conserved oligopeptide permease (Edwards et al., 2014) that was shown to influence sporulation initiation in *C. difficile*. Considering the nature of the predicted AtcS interactions, this could provide a potential mechanistic link explaining the increased biofilm formation and sporulation rate of a *hfq* mutant.

## DISCUSSION

### An online browser for easy access to the C. difficile transcriptome

In the present study, we have applied global approaches of bacterial RNA biology (Hor et al., 2018) to capture the transcriptome architecture of *C. difficile* and unravel the scope of post transcriptional regulation in this important human pathogen. Through the combination of unbiased experimental approaches to detect both transcriptional start sites and transcript 3’ ends, we have bypassed the challenge of identifying novel non-coding regulators lacking conservation to known RNA families. This is particularly important for Clostridia given their ancient origin and the significant genomic differences even between *C. difficile* strains and other pathogenic clostridia. The transcriptome maps are available for the scientific community through an open access online web browser. We intend to update our database whenever a refined genome annotation becomes available, and to extend it by transcriptome profiles obtained under growth in various infection-relevant conditions that will hopefully facilitate future genetic strategies.

### Discovery of novel regulatory functions located in 5’ UTRs

Premature transcription termination (PTT) within 5’ untranslated regions were recently shown to be widespread events in the Gram-positive bacteria *B. subtilis*, *Listeria monocytogenes* and *Enterococcus faecalis* (Dar et al., 2016). Through our transcriptome-wide mapping of 3’ transcript ends, we found 74 premature transcription termination events that were located in the 5’ UTRs of mRNA transcripts. While 57 were associated with conserved riboswitches, the remaining 17 were lacking conservation to known families of ribo-regulators. Since such PTT events often have regulatory functions, a response to unknown ligands or other unknown antitermination mechanisms is likely. Interestingly, some of the detected PTT events reside in the 5’ UTRs of multidrug-family type ABC transporters. Hence, characterizing their expression during exposure of *C. difficile* to different antibiotic molecules will potentially reveal novel mechanisms underlying antibiotic resistance in *C. difficile*. More broadly, mapping of 3 ́ ends in *C. difficile* grown under various growth conditions should reveal even more regulators that are not active during the standard laboratory growth conditions applied in this study. Such an approach recently discovered antibiotic-responsive transcription termination events upstream of ABC transporters and transcriptional regulators in *B. subtilis*, *Listeria monocytogenes* and *Enterococcus faecalis* (Dar et al., 2016) and demonstrates the widespread presence of such mechanisms as well as their potential importance for bacterial physiology.

### A large Hfq network in a Gram-positive bacterium

Our knowledge of sRNA-based gene regulation in gram-positive bacteria, particularly in *C. difficile*, is lagging behind that of gram-negative model organisms e.g. *Salmonella enterica* and *E. coli.* However, it is undisputed that sRNAs are central players in the regulation of physiological and virulence pathways in gram-positive bacteria, such as *Listeria monocytogenes*, *B. subtilis* and *Staphylococcus* species. For example, the sRNA RsaE, that is conserved in the order Bacillales, has been shown to coordinate central carbon and amino acid metabolism in *Staphylococcus aureus* (Bohn et al., 2010; Rochat et al., 2018) and also to promote biofilm formation in *Staphylococcus epidermidis* by supporting extracellular DNA release and the production of polysaccharide intercellular adhesin (PIA) (Schoenfelder et al., 2019). However, the function of Hfq in facilitating the interactions between sRNA regulators and their target mRNAs in Gram-positive bacteria remains elusive, since most of the known sRNAs seem to exert their regulatory activity independent of this global RNA-binding protein (Pitman and Cho, 2015).

Previous efforts using *in silico* applications (Chen et al., 2011) and RNA-sequencing (Soutourina et al., 2013) have predicted >100 sRNA candidates in *C. difficile*, implying the existence of a large post-transcriptional network. Here, we provide experimental evidence for the existence of 42 sRNAs (that we consider high confidence candidates) which either originate from independent transcriptional start sites or are processed from untranslated regions of mRNAs. This number is at the lower spectrum of estimated sRNAs for various bacterial species (Adams and Storz, 2020). However, considering that our analysis was done in only two different growth phases of *C. difficile* cultured in rich media, this number is likely to increase in the future. Reflecting this assumption, most of our sRNAs were expressed throughout exponential growth but decreased in stationary phase. Therefore, mapping the transcriptome under defined stress conditions will likely extend the sRNA repertoire of *C. difficile* by novel candidates that are, for example, under the control of stress-induced transcriptional regulators and therefore not expressed under the conditions that were assayed in this work.

Small regulatory RNAs are classified according to the genomic location they originate from. To date, the largest number of experimentally characterized sRNAs are those located in intergenic regions that are transcribed from their own promoter and typically have a Rho-independent terminator. However, especially sRNAs derived from the 3’ UTR of mRNAs constitute an expanding class that is gaining increasing attention as more examples are being functionally characterized (Adams and Storz, 2020; De Mets et al., 2019; Hoyos et al., 2020; Miyakoshi et al., 2019). Interestingly, only a small portion of our sRNA candidates originates from intergenic regions (8 of 42) whereas the largest class were, in fact, sRNAs derived from 3’ UTRs (18 of 42).

Many of the 3’ UTR-derived sRNAs identified in our study were associating with Hfq *in vivo* suggesting they could act in trans to potentially cross-regulate other mRNA targets. One interesting example is CDIF630nc_00006 which is generated by processing from the 3’ UTR of *cbiO*, encoding the ATP-binding protein of a cobalt-specific ABC-transporter. Cobalt is mainly incorporated into vitamin B12 which in turn is an important enzyme cofactor in a variety of biological processes in *C. difficile* (Shelton et al., 2020). A prominent one is the utilization of ethanolamine, an important carbon source for *C. difficile* during colonization of the gut whose availability impacts disease outcome (Nelson et al., 1980).

Another abundant class of sRNA (13 of 42) were *cis*-encoded antisense RNAs. Interestingly, this does not reflect genome-wide antisense transcription which only accounted for approx. 6% of TSS events in *C. difficile*. Comparative transcriptomics have revealed that bacterial antisense transcription is mainly the product of transcriptional noise and directly correlates with genomic AT content (Lloréns-Rico et al., 2016). With a GC content of only 27% in *C. difficile* this result raises the question whether our annotated antisense RNAs are functional regulators. However, the majority of *cis*-antisense encoded sRNAs (10 of 13) were bound by Hfq which suggests a regulatory function for these transcripts. If expressed in sufficient amounts they can regulate the expression of the gene encoded on the opposite strand, an interaction that is not considered to require the presence of Hfq in other studied bacteria due to extensive sequence complementarity (Irnov et al., 2010). Accordingly, we found none of these mRNAs associated with Hfq *in vivo.* This could indicate that there are additional *trans*-encoded target mRNAs regulated through these antisense sRNAs for which binding would need to be facilitated by Hfq as a consequence of imperfect complementarity.

### Implications of Hfq associations for C. difficile virulence

A recent characterization of a Hfq deletion strain revealed a hyper-sporulation phenotype of the mutant in comparison to the wild type strain suggesting that sRNAs participate in the repression of sporulation (Maikova et al., 2019). Among the Hfq-bound mRNAs we found the sporulation specific transcriptional regulator, *sigG*, and the master regulator of sporulation, *spo0A*, associated with Hfq *in vivo*. In addition, the oligopeptide transporter *oppB*, which regulates the initiation of sporulation, was enriched in the Hfq pulldown. The latter was also among the COPRA-predicted target mRNAs of the *bona fide* intergenic sRNA AtcS. Together, these mRNAs provide a first mechanistic explanation for the observed hyper-sporulation phenotype of a *hfq* mutant (Maikova et al., 2019). Until recently, only protein regulators were considered to be important for the sporulation process. However, several studies in *B. subtilis* have identified sRNA candidates to be controlled by sporulation-specific sigma factors including SigG, SigK, SigF and SigE, suggesting an involvement of RNA players in the regulation of sporulation (Marchais et al., 2011; Schmalisch et al., 2010; Silvaggi et al., 2006). Likely due to the standard growth conditions used for our experimental transcriptome annotation, we did not find any sRNA candidates that are transcribed from a sporulation-specific promoter.

Moreover, we found several mRNAs encoding for proteins involved in motility (*flgB*, *fliJ*, *flhB* and *cheW1*), the cell wall protein encoding gene *cwpV* as well as several adhesins potentially involved in biofilm formation (CDIF630_03429, CDIF630_01459 and CDIF630_03096) that could explain the observed phenotypes of a Hfq knock-down, including reduced motility, increased sensitivity to stresses and biofilm formation (Boudry et al., 2014). Some of the identified Hfq bound mRNAs, such as *spo0A* and *cwpV*, were also found to be differentially regulated in the same Hfq knock-down strain, further confirming them as direct potential targets of sRNA regulation in *C. difficile*.

In total, we identified 330 mRNAs associating with Hfq *in vivo*. This would imply that about 10% of the genome is under direct post-transcriptional regulation. However, our RIP-seq based identification of Hfq-associated mRNAs includes the sigma factors *sigA2* and the sporulation sigma factor *sigG* as well as >20 transcriptional regulators and several two component systems. Therefore, it is reasonable to assume that sRNA-based regulation could impact gene expression to a much greater extent than suggested by the 330 mRNAs enriched in our analysis.

### Identification of conserved sRNA with members outside the class Clostridia

Most of the sRNAs identified in this study are exclusively conserved within the species *C. difficile*. This lack of broader sequence conservation is probably not surprising in light of the recent re-classification of *C. difficile* that placed the species in the new taxonomic family *Peptostreptococcaceae* (Lawson et al., 2016). However, the evolution of sRNAs is far less understood than the evolution of proteins which can be conserved over long phylogenetic distances whereas individual sRNAs tend to display narrow conservation ranges restricted to the genus or family level (Jose et al., 2019). Interestingly, in some cases, sRNAs from distant phyla share the same function and target mRNAs despite a lack of primary sequence conservation. One such example is the Fur-regulated sRNA FsrA in *B. subtilis*, that regulates mRNA targets involved in iron transport and metabolism (Gaballa et al., 2008; Pi and Helmann, 2017) in a very similar way to RyhB in *Enterobacteriaceae* (Masse and Gottesman, 2002; Masse et al., 2005). The genome of *C. difficile* 630 lacks any sequence conservation to FsrA but the transcriptomic response of a *C. difficile* 630 Fur mutant involves the down-regulation of similar genes and pathways as in *B. subtilis* (Berges et al., 2018). Therefore, future analyses of transcriptional responses to iron starvation and other defined stress conditions might reveal novel sRNAs in *C. difficile* that have functionally related members in other bacteria.

Although most sRNAs were only present in *C. difficile*, two sRNAs (CDIF630nc_00069 & CDIF630nc_00001), showed a remarkably high sequence conservation outside the class Clostridia, indicating that they serve in cellular pathways that are conserved across multiple species (Frohlich et al., 2016). In Gram-positive organisms such core sRNA are not well characterized but there are a few examples, such as SR1, a dual function sRNA with broad conservation within Bacillales (Gimpel et al., 2012).

The first sRNA, CDIF630nc_00069, is associated with a putative conjugative transposon element. As a result, this sRNA is absent in some *C. difficile* strains. Hence, while CDIF630nc_00069 is present in *C. difficile* 630 (RT012) and the epidemic and hypervirulent strain R20291 (RT027), the sRNA is absent in the non-epidemic *strain* CD196 of the same RT027 ribotype (Stabler et al., 2009). Instead, CDIF630nc_00069 is sporadically found in distant bacterial lineages including *Streptococcus pneumoniae* and *Streptococcus pyogenes* (Fig. 5C and Table S10). Taken together, CDIF630nc_00069 seems to be part of a mobile genomic element that has spread amongst distant members of the phylum Firmicutes.

The second sRNA, CDIF630nc_00001, is the first experimentally validated member of the RaiA family of structured sRNA that was discovered by bioinformatic approaches (Weinberg et al., 2017), therefore, we rename it RaiA. RaiA RNAs are found in Firmicutes and Actinobacteria and share conserved nucleotides as well as secondary structures. In both phyla, they often occur upstream of the *raiA* gene that encodes for a stress-inducible ribosome-inactivating protein (Agafonov and Spirin, 2004). In *C. difficile* strains the RaiA RNA resides in the 3’ UTR of CDIF630_00250, encoding a putative phosphoribosyl transferase, whereas in other Clostridium species including *C. botulinum*, *C. perfringens*, *C. tetani* and *C. sporogenes* the sRNA is located in the commonly observed *raiA* 5’ UTR region. Our RNA-seq and RNAtag-Seq data revealed transcription of RaiA starting from a dedicated TSS and two terminators resulting in two transcripts of different lengths both of which could be detected by Northern Blot analysis (Fig. 5B).

Interestingly, neither CDIF630nc_00069 nor CDIF630nc_00001 was associating with Hfq *in vivo*. This might suggest the potential existence of alternative RNA-binding proteins in *C. difficile*, similar to *Salmonella* or *E. coli* where a second global RBP, ProQ, was recently characterized (Holmqvist and Vogel, 2018; Melamed et al., 2020; Smirnov et al., 2016). Potential candidates in Gram-positive bacteria exist, such as the KhpA/B heterodimer identified in *Streptococcus pneumoniae* which was shown to associate with several RNA classes leading to its classification as a global RNA chaperone (Zheng et al., 2017). Further, a recent Grad-seq analysis of cellular RNA-protein complexes in *Streptococcus pneumoniae* detected a KhpA/B complex that co-sedimented with tRNAs and intergenic sRNA transcripts (Hor et al., 2020). Given that we are only beginning to understand the true number and functions of sRNA and protein players involved in RNA-based gene regulation in *C. difficile*, there is great potential to discover previously unknown RNA-binding proteins in this important human pathogen.

## METHODS

### Bacterial strains and growth conditions

A complete list of *C. difficile* and *E. coli* strains that were used in this study is provided in Table S8. *C. difficile* cultures were routinely grown anaerobically inside a Coy chamber (85% N_2_, 10% H_2_ and 5% CO_2_) in Brain Heart Infusion (BHI) broth or on BHI agar plates (1.5% agar) unless indicated otherwise. When necessary, antibiotics were added to the medium at the following concentrations: thiamphenicol 15 μg/ml, cycloserine 250 μg/ml. *E. coli* cultures were propagated aerobically in Luria-Bertani (LB) broth (10 g/l tryptone, 5 g/l yeast extract, 10 g/l NaCl) or on LB agar plates (1.5% agar) supplemented with chloramphenicol (20 μg/ml) as appropriate. *E. coli* strain Top10 (Invitrogen) was used as a recipient for all cloning procedures, and *E. coli* CA434 (HB101 carrying the IncPβ conjugative plasmid R702) was used as donor strain for conjugation of plasmids into *C. difficile* 630.

### Plasmid construction

All plasmids and DNA oligonucleotides that were used are listed in Tables S9 and S10, respectively. *hfq* including its native 5’ UTR and native promoter was amplified from *C. difficile* 630 using either FFO-122 and FFO-136 or FFO-122 and FFO-207. The resulting fragments were cloned into SacI/BamHI-digested pRPF185 (Fagan and Fairweather, 2011), generating pFF-10, encoding *hfq* including 5’ UTR and native promoter, and pFF-12, encoding a C-terminally 3X-FLAG-tagged *hfq* including 5’ UTR and native promoter, respectively.

### Strain construction

pFF-10 and pFF-12 were transformed into chemically competent *E. coli* TOP10 according to standard procedures (J. Sambrook, 1989). Both strains (FFS-34, harboring pFF-10 and FFS-46, harboring pFF-12) were used for cloning and plasmid propagation. For conjugation purposes, both plasmids were transformed in *E. coli* CA434 (HB101 carrying the IncPβ conjugative plasmid R702), resulting in FFS-36 and FFS-48 respectively. Conjugation was performed according to Kirk and Fagan, 2016 (Kirk and Fagan, 2016). In short: 200 μl of *C. difficile* 630 overnight cultures were incubated at 37 °C for 2 min. Simultaneously, 1 ml of overnight *E. coli* conjugant donor culture (FFS-36 and FFS-48) was harvested by centrifugation at 4000 x g for 2 min. *E. coli* pellets were then transferred into the anaerobic workstation and gently resuspended in pre-incubated 200 μl *C. difficile* 630 culture. Following resuspension, the cell suspension was pipetted onto well-dried, non-selective BHI agar plates (10 × 10 μl spots), allowed to dry and incubated for 8 h at 37 °C. Growth was harvested using 900 μl of TY broth, serially diluted and spread on plates containing either cycloserine (control), or cycloserine and thiamphenicol, to select for transconjugants. Plates were incubated for between 24 and 72 h, until colonies were apparent. Conjugation resulted in strain FFS-38, harbouring pFF-10 and strain FFS-50, harbouring pFF-12, which were used for RIP-seq analysis.

### Total RNA extraction

Total RNA was extracted using the hot phenol protocol. Bacterial cultures were grown to the desired OD_600_, mixed with 0.2 volumes of STOP solution (95% ethanol, 5% phenol) and snap frozen at −80 °C if not directly processed. The bacterial solution was centrifuged for 20 min, 4500 rpm at 4 °C and the supernatant was completely discarded. Cells were suspended in 600 μl of 10 mg/ml lysozyme in TE buffer (pH 8.0) and incubated at 37 °C for 10 min. Next, 60 μl of 10% w/v SDS was added and everything mixed by inversion. Samples were incubated in a water bath at 64 °C, 1-2 min before adding 66 μl 3 M NaOAc, pH 5.2. Next, 750 μl acid phenol (Roti-Aqua phenol) was added, followed by incubation for 5 min. at 64 °C. Samples were briefly placed on ice to cool before centrifugation for 15 min, 13,000 rpm at 4 °C. The aqueous layer was transferred into a 2 ml phase lock gel tube (Eppendorf), 750 μl chloroform (Roth, #Y015.2) was added, and everything centrifuged for 12 min, 13,000 rpm at room temperature. For ethanol precipitation, the aqueous layer was transferred to a fresh tube, 2 volumes of 30:1 mix (EtOH:3 M NaOAc, pH 6.5) was added and incubated overnight at −20 °C. Precipitated RNA was harvested by centrifugation, washed with cold 75% v/v ethanol and air-dried. DNA contaminations were removed by DNase treatment and the RNA was re-extracted using a single phenol-chloroform extraction step. Purified RNA was resuspended in 50 µl RNase-free water and stored at −80 °C.

### Library preparation for differential RNA-seq (dRNA-seq)

Library preparation for dRNA-seq was accomplished by Vertis Biotechnology AG. In brief, total RNA was analyzed on a Shimadzu MultiNA microchip electrophoresis system. 23S/16S ratio for all samples was 1.3. RNA was fragmented via ultrasound (4 pulses a 30 sec, 4 °C) and subsequently treated with T4 Polynucleotide Kinase (NEB). Half of the samples were then treated with terminator exonuclease (TEX) for dRNA-seq.

For the cDNA synthesis, the RNA fragments were poly(A)-tailed and 5’PPP structures were removed with RNA 5’ Polyphosphatase (Epicentre). The RNA sequencing adapter with the barcodes were ligated to the 5’-monophosphate of the fragments. First strand cDNA was synthesized with an oligo(dT)-adapter primer and M-MLV reverse transcriptase. Amplification of cDNA was done via PCR to an approximate amount of 10-20 ng/µl. cDNA was purified with the Agencourt AMPure XP kit (Beckman Coulter Genomics) and analyzed with Shimadzu MultiNA microchip electrophoresis. Equimolar amounts of the samples were pooled for sequencing. cDNAs had a size between 200 and 550 bp. The library pool was fractionated via differential clean-up with the Agencourt AMPure kit. The cDNA pool was checked with capillary electrophoresis as stated above. Libraries were sequenced on an Illumina NextSeq 500 system using 75 bp read length.

### Library preparation for RNAtag-Seq protocol

Total RNA quality was checked using a 2100 Bioanalyzer with the RNA 6000 Nano kit (Agilent Technologies) and rRNA was detected. The RIN for all samples was >6.5. Equal amounts of samples (~500 ng) were used for the preparation of cDNA libraries with the RNAtag-Seq protocol as previously published by Shishkin et al. (Shishkin et al., 2015) with minor modifications.

Briefly, the RNA samples were fragmented with FastAP buffer at 94 °C for 3 min and were dephoshorylated using the FastAP enzyme (Thermo Scientific) at 37 °C for 30 min followed by bead purification with 2 volumes of Agencourt RNAClean XP beads. Fragmentation profiles were checked using RNA 6000 pico kit (Agilent) with a 2100 Bioanalyzer. The RNA fragments were ligated with 3’-barcoded adaptors at 22 °C for 1 h and 30 min using T4 RNA Ligase (NEB). Barcoded RNA samples were pooled together, the ligase was inhibited by RTL buffer (Qiagen RNeasy Min Elute Cleanup Kit) and purified with RNA Clean & Concentrator-5 column (Zymo). rRNA were depleted from pools using Ribo-Zero (Bacteria) Kit (Illumina) and purified with RNA Clean & Concentrator-5 column (Zymo). The profiles before and after rRNA depletion were analysed with RNA 6000 pico kit (Agilent) with a 2100 Bioanalyzer. The rRNA depleted RNA were reverse transcripted and the first strand of the cDNA were synthsized using a custom AR2 oligo (Sigma-Aldrich) and the AffinityScript multiple temperature cDNA synthesis kit (Agilent) at 55 °C for 55 min. The RNA was degraded with 1 M NaOH at 70 °C for 12 min, the reactions were neutralized with Acetic acid and were purified with 2 volumes of MagSi-NGS^PREP^ Plus beads (AMSBIO). A second 3Tr3 adaptor were ligated to the cDNA using T4 RNA Ligase (NEB) overnight at 22 °C followed with two bead purification steps with 2 volumes of MagSi-NGS^PREP^ Plus beads (AMSBIO). A PCR enrichment test was performed in order to determine the number of PCR cycles that are nesessery for each pool followed by a bead purification step and a QC with DNA HS kit (Agilent). Then, the final PCR was performed with the number of cycles determined from the previous step using P5 and P7 primers. Two sequential bead purification steps were performed with 1.5 and 0.7 respectively.

Libraries were quantified with the Qubit 3.0 Fluometer (ThermoFisher) and the library quality and size distribution (~480 bp peak size) was checked using a 2100 Bioanalyzer with the High Sensitivity DNA kit (Agilent). Sequencing of pooled libraries, spiked with 5% PhiX control library, was performed with ~8 million reads / sample in single-end mode on the NextSeq 500 platform (Illumina) with the High Output Kit v2.5 (75 Cycles).

We noticed a strong enrichment of reads mapping to native 3’ ends of transcripts in *C. difficile* 630 libraries that were prepared for the RNAtag-Seq protocol. We cannot fully explain this observation but we suspect a combination of inefficient RNA fragmentation and native 3’ end stability of transcripts to contribute to this outcome for RNAtag-Seq libraries prepared from *C. difficile* 630 total RNA.

### Northern blotting

RNA samples were separated on a denaturing 6% polyacrylamide gel in TBE buffer containing 7 M urea. Gels were transferred onto Hybond+ membranes (GE Healthcare Life Sciences) at 4 °C with 50 V (~100 W) for 1 h. For hybridization with P32-labeled DNA oligonucleotides, membranes were incubated over night at 42 °C in Roti^®^ Hybri-Quick Buffer (Roth). Membranes were washed three times with decreasing concentrations of SSC buffer (5 x, 1 x and 0.5 x) before imaging on a Typhoon FLA 7000 phosphor imager.

### Western blotting

To verify the expression and successful pulldown of FLAG-tagged Hfq, 9 µl lysate and 50 µl of resuspended beads were mixed with 81 µl and 50 µl 1X protein loading dye respectively and boiled for for 5 min at 98 °C. Following incubation, 20 µl of each sample was loaded and separated on a 15% SDS–polyacrylamide gel followed by transfer of proteins to a PVDF membrane. For detection of FLAG-tagged proteins, the membrane was blocked in TBS-T with 5% milk powder for 1 h at room temperature and washed 3 x in TBS-T for 10 min. Subsequently the membrane was incubated over night at 4 °C with anti-FLAG antibody (Sigma) diluted 1:1,000 in TBS-T with 3% BSA and washed again 3 x in TBS-T for 10 min. Following the last washing step the membrane was incubated for 1 h at room temperature with anti-mouse-HRP antibody (ThermoScientific) diluted 1:10,000 in TBS-T with 3% BSA and finally washed 3 x in TBS-T for 10 min before adding ECL substrate for detection of HRP activity using a CCD camera (ImageQuant, GE Healthcare).

### Hfq co-immunoprecipitation and RNA-sequencing (RIP-seq)

Overnight cultures of FFS-38 and FFS-50 were prepared in biological duplicates in buffered TY and used for inoculation of pre-cultures. Once an OD_600_ of 0.5 was reached, each replicate was used for inoculation of three separate flasks. Main cultures were grown until either mid-exponential (OD_600_=0.5), late-exponential (OD_600_=0.9) or stationary (3 h post entry) growth phase. For each condition, 50 OD were harvested by centrifugation for 20 min at 4000 rpm at 4 °C, snap frozen and resuspended in 800 µl lysis buffer (20 mM TRIS pH 8.0, 1 mM MgCl_2_, 150 mM KCl, 1 mM DTT). Subsequently, each sample was mixed with 1 µl DNase I (Fermentas, 1 U/µl) and 800 µl of 0.1 mm glass beads and lysed in a RETSCH's Mixer Mill (30 Hz, 10 min, 4°C). Bacterial lysates were cleared by centrifugation for 10 min at 14,000 x g and 4 °C. Approximately 900 µl supernatant was transferred to a new tube and incubated with 25 µl (1/2 * OD in µl) of mouse-anti-FLAG antibody (clone M2, Merck/Sigma-Aldrich #F1804) at 4 °C for 45 min (rocking). Immunoprecipitation of FLAG-tagged Hfq was performed by incubating each sample with 75 µl pre-washed (3 x resuspended in 1 ml lysis buffer and centrifuged at 10,000 rpm for 1 min) protein A sepharose beads (Merck/Sigma-Aldrich #P6649) for 45 min at 4 °C (rocking). Beads and captured proteins were washed 5 x with 500 µl lysis buffer (mixed by inversion and centrifuged at 10,000 rpm for 1 min) and resuspended in 500 µl lysis buffer. Finally, RNA, co-immunoprecipitated with antibodies and protein A sepharose beads, was eluted by adding the same volume of Phenol:Chloroform:Isoamylalcohol (25:24:1, pH 4.5, Roth). The solution was mixed for 20 s, transferred to PLG tubes (Eppendorf) and incubated at room temperature for 3 min. Following centrifugation (30 min, 15,200 x g, 15 °C) the aqueous phase was transferred to new tubes and RNA was precipitated over night at −20 °C by adding 30 µg Glycoblue and 800 µl isopropanol. Precipitated RNA was pelleted (centrifugation for 45 min, 15,200 rpm, 4 °C), washed with 80% and 100% ethanol (centrifugation for 10 min, 15,200 rpm, 4 °C) and resuspended in 15,5 µl nuclease-free water (65 °C, 1 min, 600 rpm). For DNase treatment, purified RNA was incubated with 2 µl DNase I, 0.5 µl RNase inhibitor and 2 µl 10 x DNase buffer for 30 min at 37 °C. To each sample, 100 µl nuclease free water was added and DNase treated RNA was purified by Phenol:Chloroform:Isoamylalcohol treatment, as described before. Precipitation was performed overnight at −20 °C by mixing the samples with 3 x the volume of sodium-acetate/ethanol (1:30), followed by centrifugation and washing with 80% and 100% ethanol. Finally, air-dried pellets were resuspended in 20 µl nuclease-free water (65 °C, 1 min, 600 rpm).

RNA quality was controlled using a 2100 Bioanalyzer and the RNA 6000 Pico kit (Agilent Technologies). Since rRNA was detectable, the RIN for all samples was >7.5. Equal amounts of RNA (~50 ng) were used for preparation of cDNA libraries with the NEBNext Multiplex Small RNA Library Prep kit for Illumina (NEB) with some minor changes to the manufacturers’ instructions. RNA samples were fragmented with Mg^2+^ at 94 °C for 2 min, 45 s using the NEBNext Magnesium RNA Fragmentation Module (NEB) followed by RNA purification with the Zymo RNA Clean & Concentrator kit. Fragmented RNA was dephosphorylated at the 3’ end, phosphorylated at the 5’ end and decapped using 10 U T4-PNK +/−, 40 nmol ATP and 5 U RppH respectively (NEB). After each enzymatic treatment, RNA was purified with the Zymo RNA Clean & Concentrator kit. The RNA fragments were ligated for cDNA synthesis to 3’ SR adapters and 5’ SR adapters and diluted 1:5 with nuclease-free water before use. PCR amplification was performed to add Illumina adaptors and indices to the cDNA for 16 cycles, using 1:5 diluted primers. Barcoded DNA Libraries were purified using magnetic MagSi-NGSPREP Plus beads (AMSBIO) at a 1.8 ratio of beads to sample volume. Finally, libraries were quantified with the Qubit 3.0 Fluorometer (ThermoFisher) and the library quality and size distribution (~230 bp peak size) was checked using a 2100 Bioanalyzer with the High Sensitivity DNA kit (Agilent). Sequencing of pooled libraries, spiked with 5% PhiX control library, was performed with ~5 million reads / sample in single-end mode on the NextSeq 500 platform (Illumina), using the High Output Kit v2.5 (75 cycles). Demultiplexed FASTQ files were generated with bcl2fastq2 v2.20.0.422 (Illumina). Reads were trimmed for the NEBNext adapter sequence using Cutadapt version 2.5 with default parameters. In addition, Cutadapt was given the –nextseq-trim=20 switch to handle two colour sequencing chemistry. Reads that were trimmed to length=0 were discarded.

The READemption pipeline version 4.5 (Förstner et al., 2014) with segemehl version 0.2 (Otto et al., 2014) was used for read mapping and calculation of read counts. Reads were mapped on CP010905.2 with additional annotations for ncRNAs, 3’ UTRs and 5’ UTRs. Normalization and enrichment analysis was done with the edgeR version 3.28.1 (Robinson et al., 2010). To calculate normalization factors the trimmed means of M values (TMM) method was used between flag tagged and non-flag-tagged Hfq libraries of each growth phase individually. All features that considered more than 10 read counts in at least two replicates were included in the analysis. Only features with a Benjamini-Hochberg corrected P-value of ≤0.1 and a log2 fold-change of ≥2 were considered as significantly enriched.

### Annotation of TSS, TTS and sRNAs

For the prediction of transcriptional start sites (TSSs) and small regulatory RNAs (sRNAs), we used the modular, command-line tool ANNOgesic (Yu et al., 2018). All parameters were kept at the default setting, unless stated otherwise. The TSS prediction algorithm was trained using parameters that were derived from manual curation of the first 200 kb of the genome. Secondary TSSs were excluded if a primary TSS was within seven nucleotides distance. For annotation of 5’ untranslated regions (5’ UTRs) the default settings for maximum length was corrected to 500 nt.

For the prediction of sRNAs, we used the default settings, that is, the transcript must have a predicted TSS or processing site (PS) at their 5’ end, form a stable secondary structure (calculated with RNAfold from Vienna RNA package), and a length between 30 to 500 nt. For 5’ UTR-derived sRNAs the transcripts must have a PS at their 3’ end. For 3’ UTR-derived sRNA the transcript can either have a TSS or PS at their 5’ end and share its terminator with the parental gene.

Prediction of transcriptional termination sites (TTSs) was accomplished manually. Therefore, read coverage tracks of RNAtag-Seq libraries were visualized with Integrative Genomics Viewer (Robinson et al., 2011) and scanned for enriched regions. Termination sites were defined as the first position with less than half of the maximal reads of the entire enriched region.

### Prediction of promoter and terminator motifs, novel ORFs

Motif search and generation of sequence logos was accomplished with MEME version 5.1.1 (Bailey and Elkan, 1994) and CentriMO version 5.1.1 (Bailey and Machanick, 2012). For determination of a sigA promoter motif, 100 nt upstream of each TSS was extracted and uploaded to MEME. Motifs to be found by MEME was set to 10 and only the given strand was searched. All other settings were left at default. The resulting sigA motif matrix was then uploaded to CentriMo along with the extracted promoter regions. Again, only the given strand was searched and all other settings were left at default. Based on the CentriMo output, those promoter sequences harboring a sigA promoter motif were uploaded to MEME ones more to refine the sigA promoter motif. In contrast to sigA, previously published promoter motifs harboring either a sigB, sigH or sigK promoter motifs were uploaded for the initial MEME search instead of extracted promoter sequences generated in this study. The input sequences were based on the following studies: sigB (Kint et al., 2017); sigH (Saujet et al., 2011); sigK (Saujet et al., 2013). The thereby generated motif matrices were uploaded to CentriMo together with our extracted promoter sequences. Those sequences harboring a sigB, sigH or sigK motif were uploaded to MEME to generate a refined promoter motif based on our input sequences. Following the MEME search, some sequences were excluded manually based on the following criteria: promoter location within the uploaded sequence has to be at the 3’ end; at least one G within 3 nucleotides next to the −10 box (only sigB); G at position 25, 26 or 27 (only sigH); ATA at position 23-25 and at least one A and C between position 2-7 (only sigK). Finally, the manually curated promoter sequences were uploaded to MEME to generate the final promoter motif.

For 3’ UTR motif search 15 nt downstream of all termination sites were extracted to search for the presence of the common poly-U stretch. Command line version of MEME was executed with default options except for the minimal motif length, which was set to 4. Only the given strand was used for the search.

Prediction of novel ORFs was accomplished manually. In order to identify novel ORFs we focused on oTSS, - TSS not associated with any coding or non-coding sequence. To avoid any miss-annotation we applied the following criteria: (i) presence of a start and stop codon (ii) presence of a ribosome binding site within 15 bp upstream of a start codon (iii) sequence conservation in other *Clostridioides* strains.

### Prediction of RNA folding and sRNA target mRNAs

Secondary structures of ncRNAs were predicted with the RNAfold WebServer (Lorenz et al., 2011) and visualized with VARNA (Darty et al., 2009). To predict folding of TTS the sequence 60 nt upstream and 10 nt downstream of the termination site was extracted. RNAfold version 2.2.7 of the Vienna RNA Package 2 was used with default settings to predict secondary structures.

Potential sRNA-target interactions were predicted using CopraRNA (Raden et al., 2018; Wright et al., 2014; Wright et al., 2013). Target prediction was performed for all sRNAs that had an associated Hfq peak in our RIP-seq dataset. As input, three homologous sRNA sequences were uploaded for each sRNA including the *C. difficile* 630 homologue. Due to the low sequence conservation, only homologues present in other *C. difficile* strains could be selected as input sequences, however, we focused on those sequences that showed at least some sequence variation when compared to *C. difficile* 630. The target region was specified as 200 nt upstream and 100 nt downstream of annotated start codons. Default settings were used. CopraRNA predictions for each sRNA candidate were then compared with mRNA candidates that co-immunoprecipitated with Hfq to filter for high probability targets.

## DATA AVAILABILITY

All RNA-sequencing data are available at the NCBI GEO database (http://www.ncbi.nlm.nih.gov/geo) under the accession number GSE155167.

## ACKNOWLEDGEMENT

We thank the Core Unit SysMed at the University of Würzburg for technical support with RIP seq and RNAtag-Seq data generation. This work was supported by the IZKF at the University of Würzburg (project Z-6). We thank the Vogel Stiftung Dr. Eckernkamp for supporting F.P. with a Dr. Eckernkamp Fellowship.

## AUTHOR CONTRIBUTIONS

MF, VL-S, JV, and FF conceived and designed the study. MF and V-LS performed experiments, LJ, V-LS and LB setup the web browser. VL-S, FP and LB analyzed RNA-seq data. FF supervised the project. MF, VL-S and FF wrote the manuscript.

