## Supplemental figures S1-2; Supplemental tables S9-10 for "An RNA-centric global view of *Clostridioides difficile* reveals broad activity of Hfq in a clinically important Gram-positive bacterium"

**Figure S1** Conservation analysis of *C. difficile* 630 sRNAs. Conservation of sRNAs was analysed with profile Hidden Markov Models against a genome database of Refseq assemblies of the phylum Firmicutes. Percent identities of 3UTR, 5UTR, IGR and cis-antisense ncRNA against selected strains are shown. For several hits in one genome, only the hit with the highest percent identity is shown. Rows and columns were hierarchically clustered with UPGMA (unweighted pair group method with arithmetic mean) algorithm.

**Figure S2** The spectrum of Hfq-associated sRNAs and mRNAs in *C. difficile* 630 across different growth phases. (A) Distribution of reads matching experimentally annotated sRNA candidates in Hfq-FLAG cDNA libraries at different phases of growth. Percentage indicates the reads (in transcripts per million) of a given sRNA compared to all sRNAs in a cDNA library. (B) Scatter-plot analysis of RIP-seq results for mRNAs that were enriched in stationary phase (log2 f.c. ≥2; cDNA read ≥10; Benjamini-Hochberg corrected P-value ≤0.1) in the 3xFLAG tagged Hfq samples. Genes encoding for virulence-related proteins are labelled.

### Figure S1

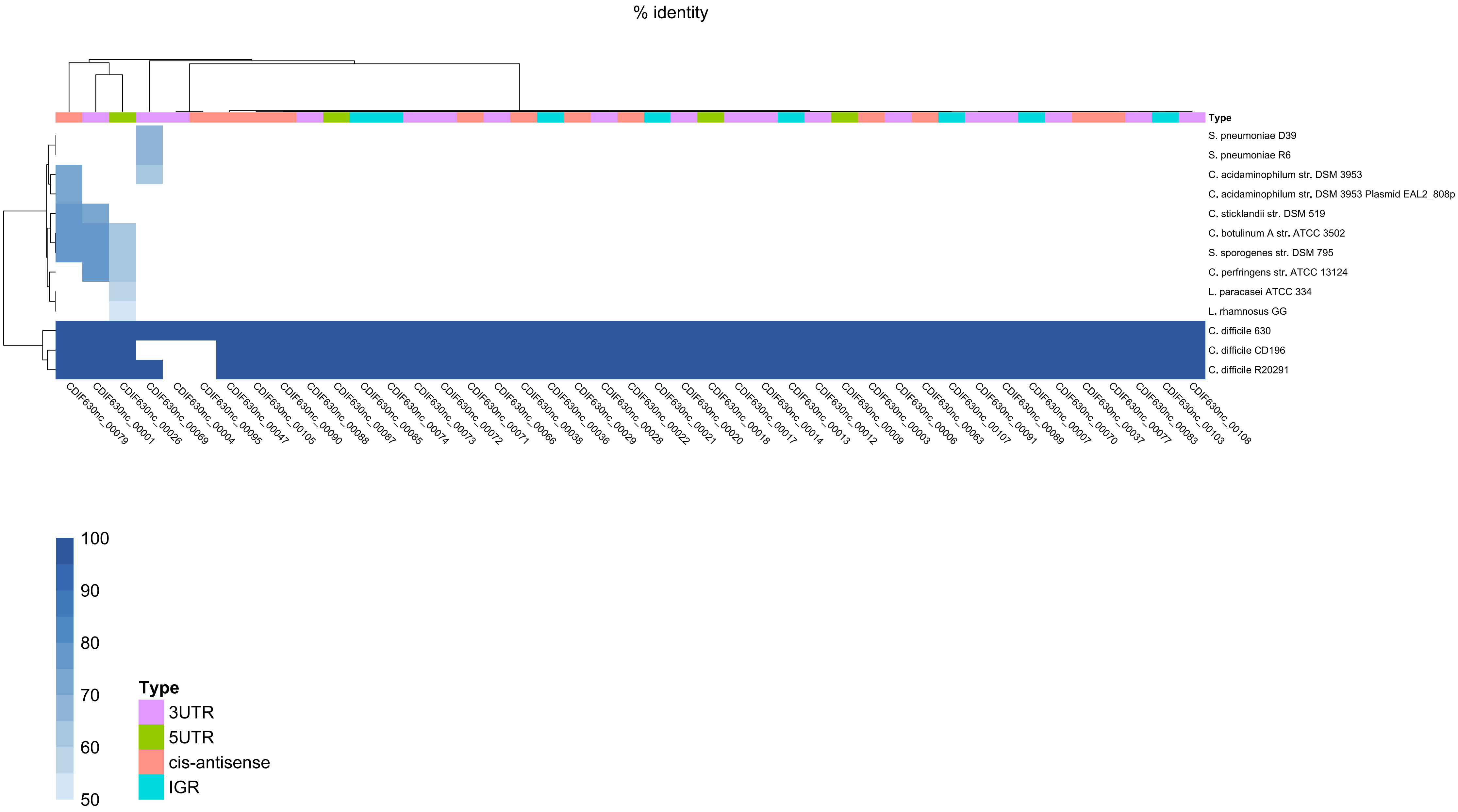

% identity
Type
S. pneumoniae D39
S. pneumoniae R6
C. acidaminophilum str. DSM 3953
C. acidaminophilum str. DSM 3953 Plasmid EAL2_808p
C. sticklandii str. DSM 519
C. botulinum A str. ATCC 3502
S. sporogenes str. DSM 795
C. perfringens str. ATCC 13124
L. paracasei ATCC 334
L. rhamnosus GG
C. difficile 630
C. difficile CD196
C. difficile R20291
CDIF630nc_00079
CDIF630nc_00001
CDIF630nc_00026
CDIF630nc_00069
CDIF630nc_00004
CDIF630nc_00095
CDIF630nc_00047
CDIF630nc_00105
CDIF630nc_00090
CDIF630nc_00088
CDIF630nc_00087
CDIF630nc_00085
CDIF630nc_00074
CDIF630nc_00073
CDIF630nc_00072
CDIF630nc_00071
CDIF630nc_00066
CDIF630nc_00038
CDIF630nc_00036
CDIF630nc_00029
CDIF630nc_00028
CDIF630nc_00022
CDIF630nc_00021
CDIF630nc_00020
CDIF630nc_00018
CDIF630nc_00017
CDIF630nc_00014
CDIF630nc_00013
CDIF630nc_00012
CDIF630nc_00009
CDIF630nc_00003
CDIF630nc_00006
CDIF630nc_00063
CDIF630nc_00107
CDIF630nc_00091
CDIF630nc_00089
CDIF630nc_00007
CDIF630nc_00070
CDIF630nc_00037
CDIF630nc_00077
CDIF630nc_00083
CDIF630nc_00103
CDIF630nc_00108
100
90
80
70
60
50
Type
3UTR
5UTR
cis-antisense
IGR

**Figure S2**

**A**

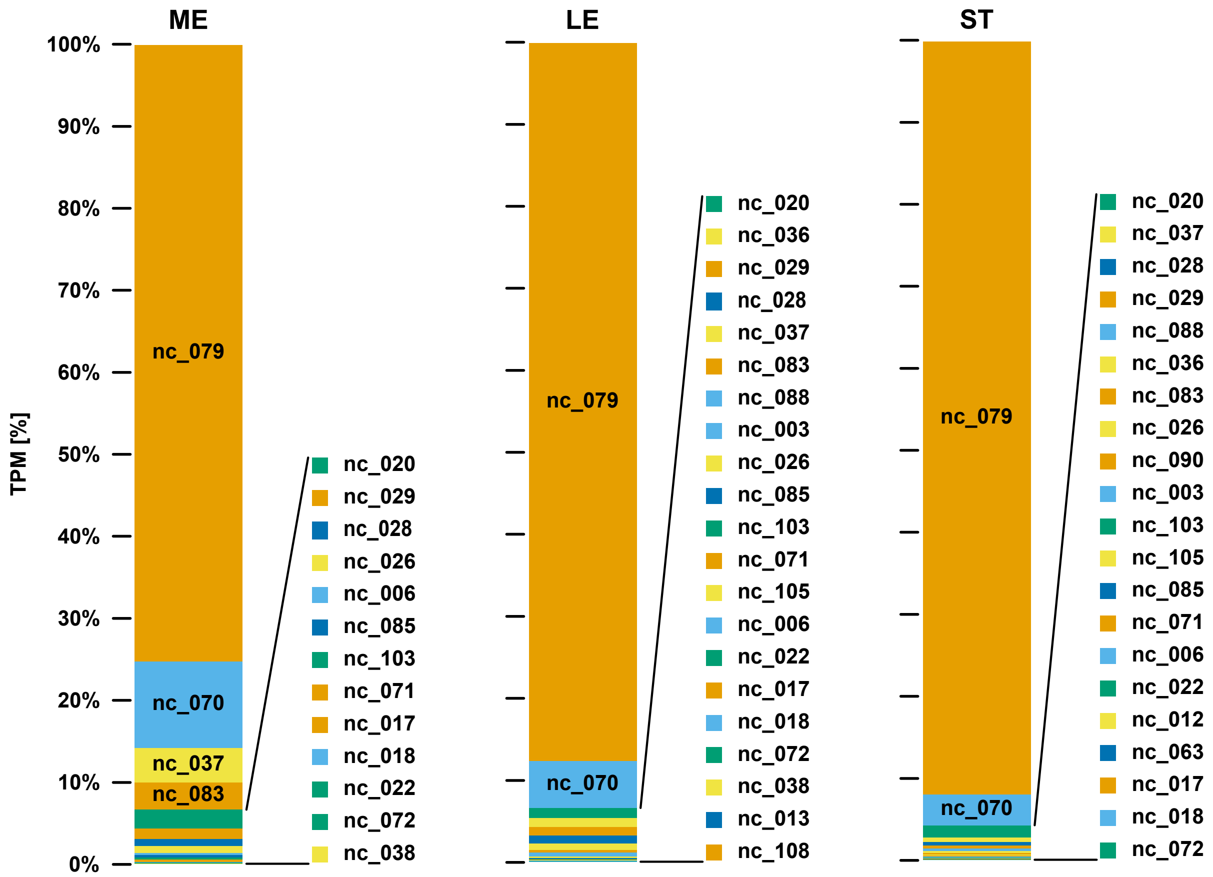
ME
LE
ST
TPM [%]
100%
90%
80%
70%
60%
50%
40%
30%
20%
10%
0%
nc_079
nc_070
nc_037
nc_083
nc_020
nc_029
nc_028
nc_026
nc_006
nc_085
nc_103
nc_071
nc_017
nc_018
nc_022
nc_072
nc_038
nc_036
nc_083
nc_088
nc_003
nc_105
nc_013
nc_108
nc_090
nc_012
nc_063

**B**

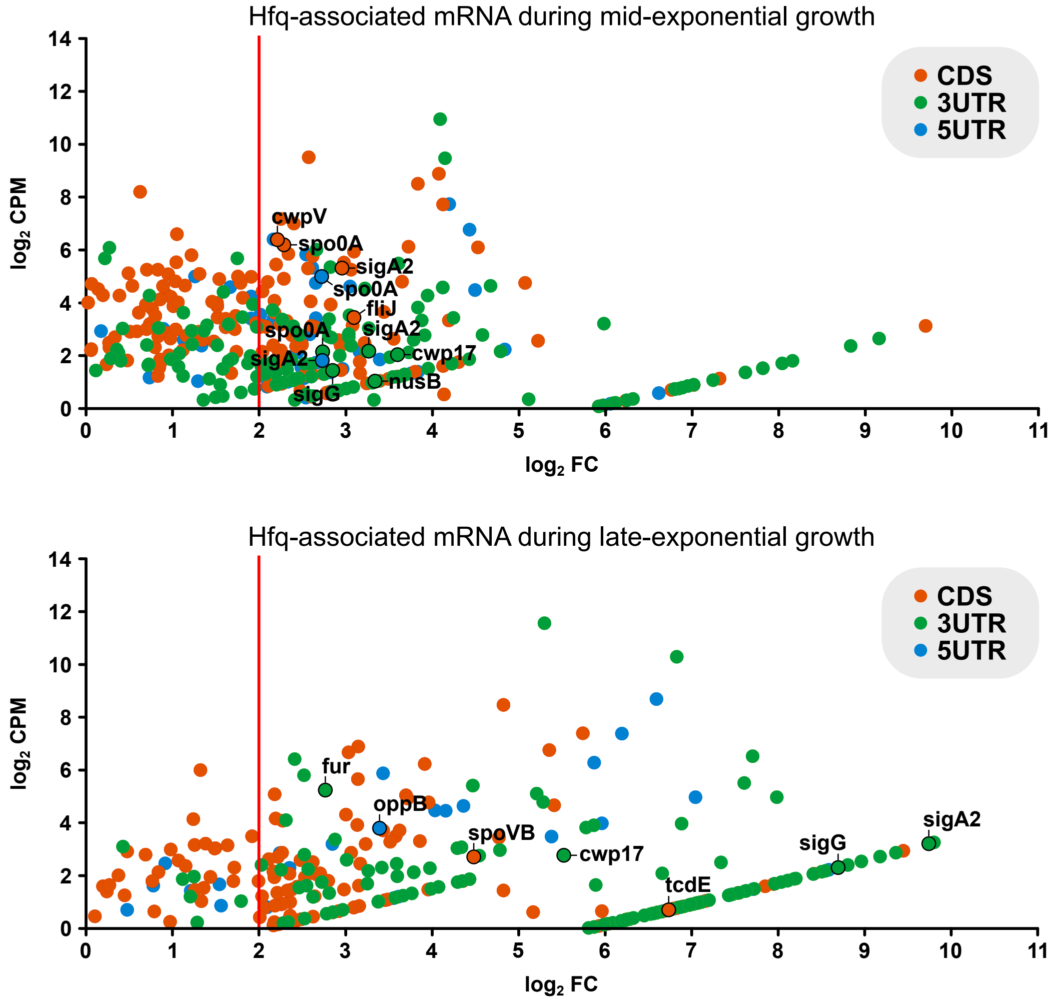
Hfq-associated mRNA during mid-exponential growth
CDS
3UTR
5UTR
log2 CPM
log2 FC
cwpV
spo0A
sigA2
spo0A
fliJ
spo0A
sigA2
cwp17
sigA2
nusB
sigG
Hfq-associated mRNA during late-exponential growth
CDS
3UTR
5UTR
log2 CPM
log2 FC
fur
oppB
spoVB
cwp17
sigG
sigA2
tcdE

*Table S8: Bacterial strains used in this study*

| Strain | Relevant markers / Genotype | Origin |
|---|---|---|
| *Escherichia coli* | | |
| TOP10 | F- mcrA Δ(mrr-hsdRMS-mcrBC) φ80lacZΔM15 ΔlacX74 nupG recA1 araD139 Δ(ara-leu)7697 galE15 galK16 rpsL(StrR) endA1 λ- | Invitrogen |
| CA434 | thi-1 hsdS20 (r-B, m-B) supE44 recAB ara-14 leuB5proA2 lacY1 galK rpsL20 (strR) xyl-5 mtl-1 | Dieter Jahn |
| FFS-34 | Top 10 carrying pFF-10 | this study |
| FFS-36 | CA434 carrying pFF-10 | this study |
| FFS-46 | Top 10 carrying pFF-12 | this study |
| FFS-48 | CA434 carrying pFF-12 | this study |
| *Clostridioides difficile* | | |
| 630 | 630 wild-type strain | DSMZ |
| FFS-38 | 630 carrying pFF-10 | this study |
| FFS-50 | 630 carrying pFF-12 | this study |

*Table S9: Plasmids used in this study.*

| Plasmid | Relevant details | Origin |
|---|---|---|
| pRPF185 | *C. difficil*e inducible expression system. Shuttle plasmid containing a tetracycline-inducible gusA | Dieter Jahn |
| pFF-10 | Plasmid for expression of Hfq from its native promotor | this study |
| pFF-12 | Plasmid for expression of C-terminally 3XFLAG tagged Hfq from its native promotor | this study |

*Table S10: DNA oligonucleotides used in this study.*

| Oligo | Sequence (5'-3') | Description |
|---|---|---|
| *Plasmid construction* | | |
| FFO-122 | GTTAACAGATCTGAGCTTATACAACTTAATATTGAAAATTTGTC | amplification of *hfq* including native promoter for Gibson cloning into pRPF185 |
| FFO-136 | AAGTTTTATTAAAACTTATAGCTATCTGTTGTTATTATTATTGTTGTTTTG | amplification of *hfq* including native promoter for Gibson cloning into pRPF185 |
| FFO-207 | AAGTTTTATTAAAACTTATAGCTACTTGTCATCGTCATCCTTGTAGTCGATGTCATGATCTTTATAATCACCGTCATGGTCTTTGTAGTCTCTGTTGTTATTATTATTGTTGTTTTG | amplification of *hfq* including native promoter and C-terminal 3XFLAG-tag for Gibson cloning into pRPF185 |
| *Northern blot probes* | | |
| FFO-28 | GCTTCATCAATTTTTCCGTA | targeting CDIF630nc_00010 |
| FFO-34 | CGTAAGGTGAGAAGCGGACT | targeting CDIF630nc_00018 |
| FFO-35 | ACTAGGGTTACCAGGGGGAT | targeting CDIF630nc_00021 |
| FFO-49 | ATCAAATATCACCGAACAAC | targeting speF RSW |
| FFO-56 | CCAACTATAGCAATACCTCA | targeting CDIF630nc_00079 |
| FFO-60 | AGTTCATTTTGAGACACCCT | targeting CDIF630nc_00089 |
| FFO-61 | AGCTTCTTTTTCTGTGTGAG | targeting CDIF630nc_00090 |
| FFO-67 | CTGACCTGGTTTTTGACCCCACACTCGCCT | targeting CDIF630nc_00001 (RaiA) |
| FFO-69 | TCCTGCCTGCATTACCAGTAACATGTCTTC | targeting CDIF630nc_00008 |
| FFO-87 | GATTTGGCAGGGCGGCGTATCCTGCATCTC | targeting CDIF630nc_00070 |
| FFO-211 | TCAGAAAACCATAGCCTTCTGAAACAACTT | targeting CDIF630nc_00085 (AtcS) |
| FFO-228 | ATAGCACCTATTGGCGTAGGTACTATACTA | targeting CDIF630nc_00022 |
| FFO-230 | AGTTTGAAATTAGTTAACTAACTTGCTGAT | targeting CDIF630nc_00009 |
| FFO-233 | GAAGGTAGCGGGGGATTTCCGACTACCTTC | targeting CDIF630nc_00028 |
| FFO-237 | TTCTCATTGTCCTAGTTACCTTGCCCTTTC | targeting CDIF630nc_00084 |
| FFO-238 | TAACCTAGGTATTGCCTGCTTACCAGTAAC | targeting CDIF630nc_00086 |
| FFO-239 | CACACATATAGTCGCGCAAATACTCTATCT | targeting CDIF630nc_00087 |
| FFO-293 | CCTTATGATTTGGGGAATCATAAGGCAC | targeting CDIF630nc_00006 |
